# Stem cell control and cancer initiation by an autocrine, injury-activated Igf complex

**DOI:** 10.64898/2026.02.02.703150

**Authors:** Yue Zhang, Youcef Ouadah, Yin Liu, Maya E. Kumar, Makenna M. Morck, Mark A. Krasnow

## Abstract

Stem cells rapidly proliferate after injury to repair damaged tissue, and chronic injury predisposes to cancer. However, injury-activated mitogens, the mechanisms that keep them inactive until injury, and their role in cancer are not understood. Here we identify Igf2 as the injury-activated mitogen for neuroendocrine stem cells, a facultative airway stem cell and origin of small cell lung cancer. Igf2 is constitutively produced by the stem cells but sequestered in inactive form by co-expressed Igf binding proteins. Injury releases Igf2 and induces proliferation by activating its receptors and repressing Rb tumor suppressor, which normally enforces stem cell quiescence. Persistent pathway activation initiates oncogenesis. Thus, in addition to its classical hormonal roles in physiology, growth, and aging, Igf operates locally with Igf binding proteins and Rb to control injury-induced stem cell activation and cancer. This pathway may also control related stem cells and cancers of the body and brain.

## Introduction

Adult stem cells are rapidly but transiently induced to proliferate after injury to repair damaged tissue (*1*), and repeated or chronic injury greatly increases the risk of cancer (*2, 3*). Although there has been good progress in identifying and characterizing mammalian stem cells (*4–6*) and homeostatic mitogens that regulate their proliferation (*7–10*), injury-induced mitogens, their regulatory mechanisms, and if and how they contribute to cancer (*11–13*) have remained elusive. Here we describe the long sought mitogen for a lung stem cell, its injury-activated regulation and effector program (*14–17*), and role in initiating small cell lung cancer (SCLC) (*16–21*). Surprisingly, the mitogen is a classical hormone that here functions instead as an autocrine stem cell ligand that is sequestered in the niche by a co-expressed inhibitor until liberated by injury, then acts through a canonical tumor suppressor to trigger the first step in the stem cell program and cancer initiation. The results have important implications for detecting and controlling the pre-malignant stage of one of the deadliest cancers, and for related stem cells and cancers across the body and brain.

## Results

### Igf ligands induce neuroendocrine cell proliferation in mouse lung slice culture

Pulmonary neuroendocrine (NE) cells are rare lung epithelial cells with specialized sensory, secretory, and stem cell functions (*22–27*). Many are found in clusters of 20-30 cells (neuroendocrine bodies, NEBs) that monitor airway oxygen, chemicals, and mechanical deformation, and signal this sensory information locally in the lung, to the brain through synapses with pulmonary sensory neurons, and potentially globally through secretion of myriad neurotransmitters and neuropeptides and hormones. A subset of NE cells, termed NE^stem^, function as facultative stem cells, proliferating extensively after airway injury to form large clonal repair patches that restore the surrounding epithelium (*16, 17, 28, 29*). These stem cells can also be activated in the absence of injury by compound deletion of *Rb* and *p53* (mouse *Rb1*, *Trp53;* human *RB1*, *TP53*) (*16, 18, 19*), the two defining tumor suppressors compromised in SCLC (*30, 31*). In this case stem cell activation is permanent and initiates SCLC.

To identify the mitogenic pathway for NE^stem^, we examined expression in adult mouse pulmonary NE cells (*17*) of 77 receptors or receptor subunits curated for functions in cell proliferation (*32*) (e.g., receptor tyrosine kinases), stem cell control (*33, 34*) (e.g., receptors in the Hedgehog and Wnt pathways), or SCLC (*21*) (e.g., opioid and nicotinic acetylcholine receptors) (Fig. 1A, table S1). This identified 40 expressed receptors (>10 TPM, range 1-82% of NE cells) (Fig. 1B, table S1), among which epidermal growth factor receptor (*Egfr*) was as an appealing candidate given its role as a mitogenic receptor for many epithelial stem cells (*35–37*) including in lung (*38–43*). However, pan-epithelial deletion of *Egfr* (using *Shh^EGFP-Cre^* (*44*) and *Egfr^flox^* (*45*)), did not prevent NEB development (fig. S1A) or the proliferative response of NE^stem^ in adults to airway injury induced by naphthalene, as tracked by incorporation of synthetic nucleoside EdU (*46*) (fig. S1B-E, table S2 contains statistical information related to Figures throughout).

**Figure 1.**
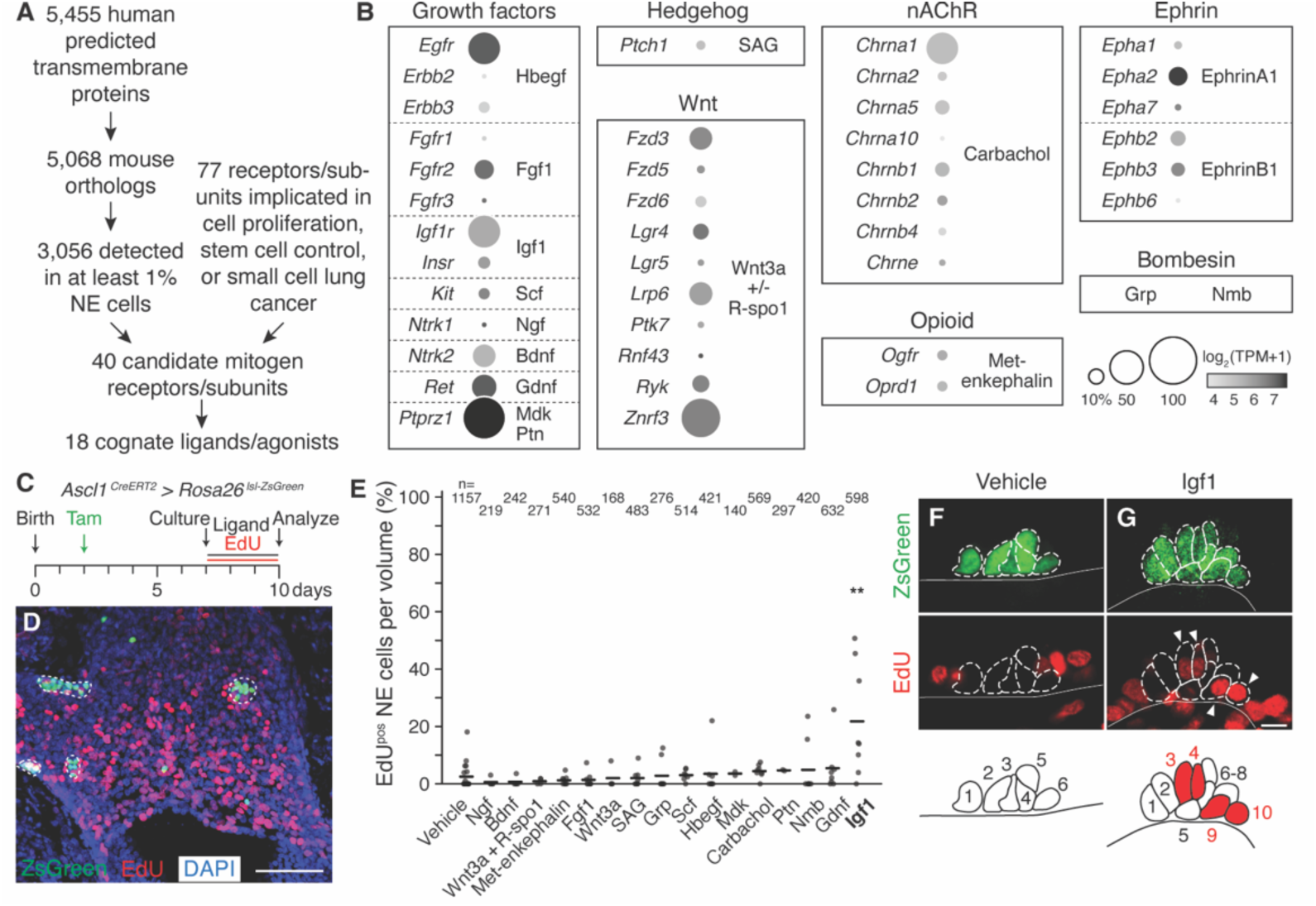
Screen for neuroendocrine (NE) cell mitogens in mouse lung slice culture. **(A)** Computational filtering scheme used to identify candidate mitogen receptors expressed in mouse pulmonary NE cells from scRNA-seq profiles (*17*) and the cognate ligands or agonists tested. “Mitogen” is the NE^stem^ mitogenic signal that induces early self-renewal after airway injury. **(B)** Mean expression level (dot heatmap, log_2_(TPM+1)) and fraction of expressing pulmonary NE cells (dot size, percent NE cells TPM>10) of the candidate receptor or receptor subunit genes indicated (grouped by signaling family) in the pulmonary NE cell scRNA-seq dataset (*17*). Bombesin family receptors (Grpr, Nmbr) are not detected in NE cells in the scRNA-seq dataset analyzed but their ligands (Grp, Nmb) were included here because they are expressed by some pulmonary NE cells. TPM, transcripts per million mapped reads. **(C)** Experimental scheme to test mitogenic effects of ligands in mouse lung slice culture. NE cells were labeled with ZsGreen by administering tamoxifen (Tam) to *Ascl1^CreERT2/+^; Rosa26^lsl-ZsGreen/+^* pups at P2, and five days later (P7) lungs were isolated and bronchial slice cultures initiated. Ligands along with EdU to track cell division were supplied in the culture medium throughout the full culture period (72 hours). **(D)** Maximum intensity projection of a bronchial slice z-stack that was stained for EdU (red) with DAPI nuclear counterstain (blue) following culture. NE cells are detected by ZsGreen native fluorescence (green). Dotted outlines, NE cell clusters (neuroendocrine bodies, NEBs). Other (non-NE) epithelial cells in the culture are often detected proliferating. Scale bar, 100 μm. **(E)** Fraction of NE cells in each slice volume imaged that proliferated during the culture period (EdU^pos^) in vehicle control and during treatment with each indicated ligand. Thick horizontal lines, distribution means (sample sizes, mean values, and additional details are given in table S2 and Methods). Only Igf1 induced significant NE proliferation (mean 21.8% EdU^pos^ vs. 2.5% in vehicle control, Cohen’s *d* 1.75, p<0.01). n, total number of NE cells scored. **(F, G)** Photomicrographs of representative NEBs quantified in (E). Note no NE cells (ZsGreen^pos^, dashed outlines) were EdU^pos^ (red) in NEB from vehicle control slice culture (F), but two pairs of NE cells were EdU^pos^ in NEB from Igf1-treated culture (G, white arrowheads). Schematics below show NE cells outlined and numbered, and EdU^pos^ NE cells filled red; black line (white line in micrographs), bronchial epithelium basement membrane. Some surrounding (non-NE) cells proliferated (EdU^pos^) in both cultures. Scale bars, 10 μm.

We then tested cognate ligands or agonists of each of the expressed candidate receptors or receptor families (Fig. 1B, table S1) for the ability to induce NE proliferation, using mouse postnatal lung slice cultures with genetically labeled NE cells (*Ascl1^CreERT2^* (*17, 47–50*) > ZsGreen (*51*)), and EdU added to the culture medium to mark cells that proliferate (Fig. 1C, D). Among the 18 ligands tested, only Igf1 induced a substantial (8.7-fold) and statistically significant increase in NE proliferation (21.8% EdU^pos^ NE cells vs. 2.5% in vehicle controls, Cohen’s *d* 1.75, p<0.01 Dunn’s test) (Fig. 1E-G, fig. S2, table S1). The EdU^pos^ NE cells were typically found as pairs, suggesting newborn daughter cells did not migrate far from their parent, but without clear stereotypy in the position of proliferative NEBs (e.g., nodal vs. internodal) or of proliferative cells within each NEB (e.g., apical vs. basal, central vs. peripheral) (Fig. 1G). The proportion of EdU^pos^ NE cells and their scattered positioning after Igf1 treatment matched results obtained after acute airway injury (10-20% NE cells proliferate) (*15–17*), suggesting that the added Igf acts selectively and efficiently on NE^stem^ in the cultures. Igf2 also induced proliferation and with similar efficiency as Igf1 (Fig. 3F).

### Igf receptors are required for injury-induced proliferation of NE^stem^

Igf1 and Igf2 exert their signaling effects primarily through Igf1r (*52*) (Igf1 receptor). In two scRNA-seq datasets, *Igf1r* is expressed (TPM>10) in 27% (*53*) and 50% (*17*) of pulmonary NE cells (n=180, 100 NE cells, respectively) (Fig. 1B, fig. S3A, table S1), and similar results obtained by single molecule fluorescence *in situ* hybridization (smFISH, 39% NE cells) (fig. S4C, left bar). To investigate involvement of *Igf1r-*expressing NE cells in the proliferative response to airway injury, we first examined EdU incorporation following naphthalene challenge *in vivo* (fig. S4A). Without injury, *Igf1r^pos^*NE cells were uniformly EdU^neg^, consistent with extremely limited or absent NE stem cell activity under homeostatic conditions (*15–17, 54*) (fig. S4C, left bar). Following injury, 43% of *Igf1r^pos^* NE cells were EdU^pos^, indicating that at least some *Igf1r^pos^* cells function as NE^stem^ after injury (fig. S4B, C). Despite their proliferation, the fraction of *Igf1r^pos^*NE cells increased little after injury (43% vs. 39%, p=0.39 binomial test), suggesting that some daughter cells turn off the receptor. Indeed, about half of the EdU^pos^ NE cells (45%) did not detectably express *Igf1r*, the expected result if following each NE^stem^ division one daughter cell retains expression while the other loses it.

To determine if NE^stem^ require Igf1r to proliferate after injury, we deleted the receptor from NE cells using *Ascl1^CreERT2^*and a conditional loss-of-function *Igf1r* allele (*Igf1r^flox^*(*55*)), while also marking the manipulated cells with ZsGreen. Two weeks after *Igf1r* deletion in adults, mice were treated with naphthalene and NE proliferation was tracked with EdU again for one week (Fig. 2A). NE *Igf1r* deletion reduced EdU incorporation by 62%, from 26% of NE cells in wild type to 10% in *Igf1r* homozygous mutants (Cohen’s *d* −0.88, p<10^−6^ Dunn’s test), whereas heterozygous *Igf1r* deletion had no effect (Cohen’s *d* −0.25, p=0.26 Dunn’s test) (Fig. 2B-D). Igf1r signaling is therefore required cell-autonomously for NE^stem^ proliferation after injury.

**Figure 2.**
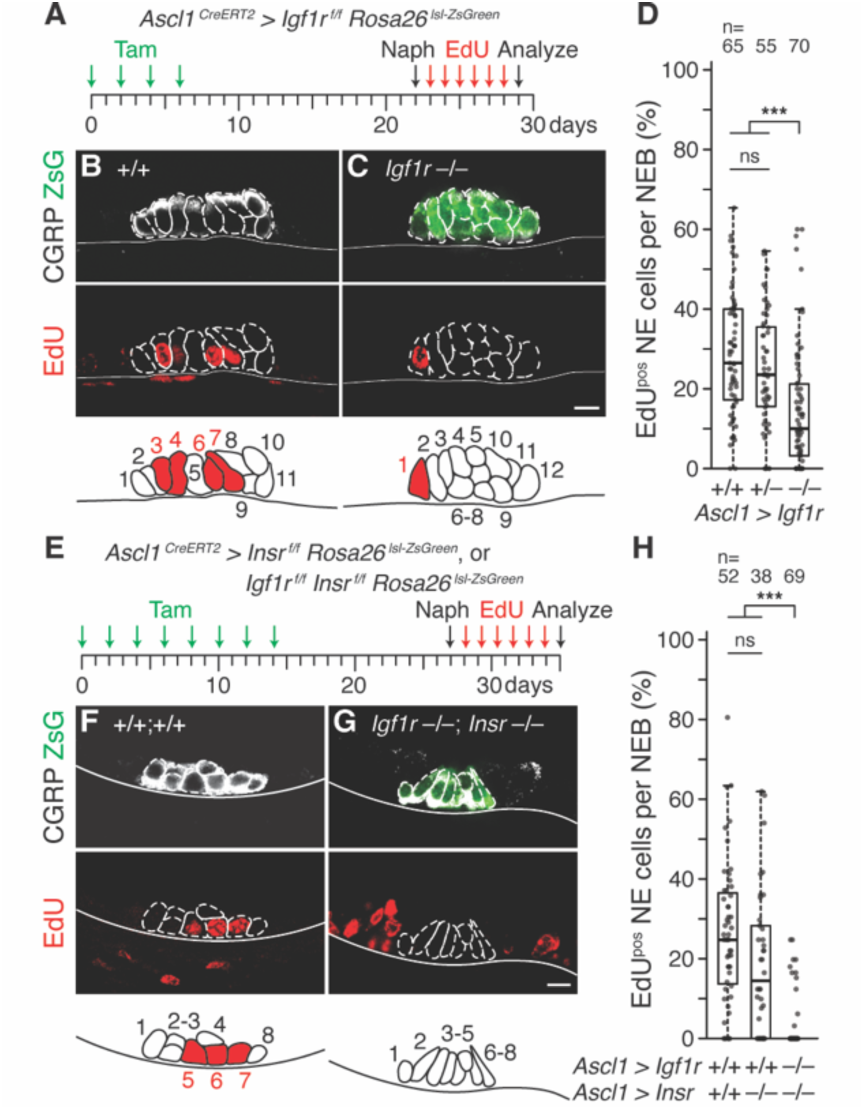
Effect of Igf1 and insulin receptor conditional deletion on NE^stem^ proliferation after airway injury. **(A)** Scheme for testing *Igf1r* requirement for NE^stem^ proliferation after airway injury with naphthalene. Adult *Igf1r^flox/flox^ Rosa26^lsl-ZsGreen/ZsGreen^* (+/+), *Ascl1^CreERT2/+^ Igf1r^flox/+^ Rosa26^lsl-ZsGreen/ZsGreen^* (+/–) or *Ascl1^CreERT2/+^ Igf1r^flox/flox^ Rosa26^lsl-ZsGreen/ZsGreen^* (–/–) mice were induced with tamoxifen (every other day, 4 total doses) to excise an essential *Igf1r* exon (and label NE cells with ZsGreen). After 16 days for recombination and Igf1r protein turnover, naphthalene (Naph) was administered and daily EdU injections tracked cumulative proliferation *in vivo* for one week after injury. **(B, C)** Photomicrographs and NEB schematics of control *Igf1r* wild type (littermate lacking *Ascl1^CreERT2^*driver allele, panel B) and NE *Igf1r* conditional deletion (C) NEBs immunostained for CGRP (NE marker, white) and EdU (red) one week after injury. Note fewer NE cells proliferated (EdU^pos^) in *Igf1r* conditional deletion. ZsGreen signal in (C) is an additional, genetic NE marker due to presence of *Ascl1^CreERT2^*driver that is missing in littermate control (B). Scale bars, 10 μm. **(D)** Quantification of (B, C). Boxplots showing the fraction of NE cells per NEB that proliferated in each NE *Igf1r* genotype. Thick horizontal lines, distribution medians. Conditional homozygous *Igf1r* deletion reduced NE proliferation by 62% (–/– vs. +/+, Cohen’s *d* −0.88, p<10^−6^), whereas conditional heterozygous deletion had no significant effect (+/– vs. +/+, Cohen’s *d* −0.25, p=0.26). n, number of NEBs scored; ns, not significant. **(E)** Scheme for testing *Insr* alone and *Igf1r* and *Insr* compound requirement. As in panel (A) except an *Insr^flox/flox^* allele was crossed in and additional doses of tamoxifen were used to facilitate compound deletion of the *Igf1r* and *Insr* conditional alleles. **(F, G)** Photomicrographs and NEB schematics as above one week after injury. Note no NE cells proliferated (EdU^pos^) in the compound *Igf1r Insr* conditional deletion (G). Scale bars, 10 μm. **(H)** Quantification of (F, G). *Insr* only deletion has no significant effect (Cohen’s *d* 0.26, p=0.33). *Igf1r Insr* compound deletion reduced NE proliferation by 87% (mean, Cohen’s *d* −1.62, p<10^−5^). n, NEBs scored.

**Figure 3.**
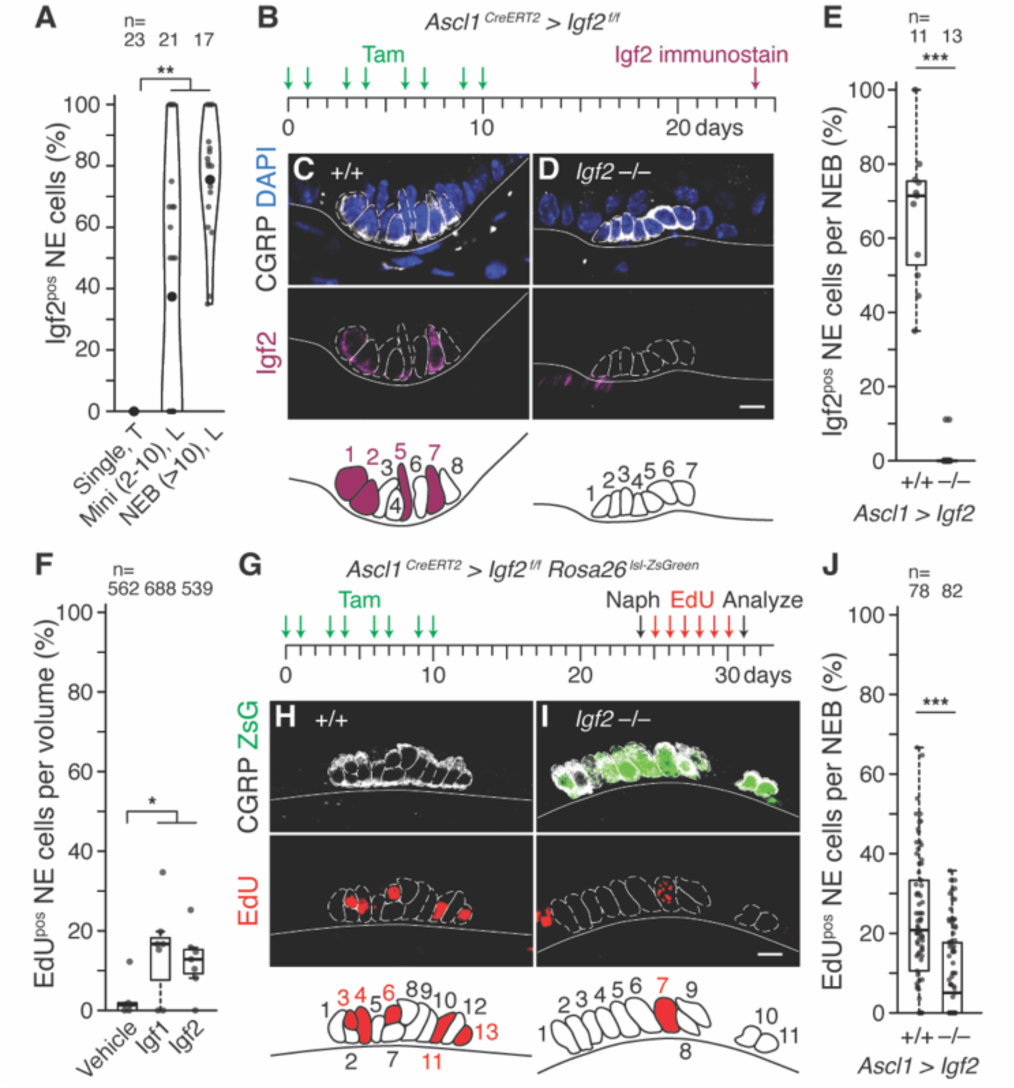
Igf2 expression and requirement in NEBs for NE^stem^ proliferation after airway injury. **(A)** Violin plots showing fraction of Igf2^pos^ NE cells detected by immunostaining for Igf2 and CGRP with DAPI counterstain in mouse trachea (“T”, all solitary NE cells) and intrapulmonary (“L”) mini-clusters (Mini, 2-10 NE cells/cluster) and NEBs (>10 NE cells/cluster). fig. S5A, B, I are representative images used for quantification. n, number of each type scored; small dots, one NE cluster or singleton; large dots, mean; Cohen’s *d* −1.38 – −1.19, p<10^−2^. Among mini-clusters with at least one Igf2^pos^ cell, three-quarters of NE cells were Igf2^pos^, similar to the value for NEBs. **(B)** Scheme for assessing efficiency of *Igf2* conditional deletion from NE cells. Adult *Ascl1^CreERT2/+^; Igf2^flox/flox^* mice were treated with tamoxifen to excise an essential exon of *Igf2*, and NEBs were analyzed by immunostaining as in (A) two weeks later. **(C, D)** Igf2 immunostaining and schematics of representative NEBs without (C, *Igf2^flox/flox^*, +/+) and with *Igf2* conditional deletion (D, *Ascl1^CreERT2/+^ Igf2^flox/flox^*, –/–). Samples were co-stained for CGRP (white, NE cells) with DAPI counterstain (blue). Igf2 protein is detected in half of the visible NE cells in (C) but none in (D). Scale bars, 10 μm. **(E)** Quantification of (B-D) showing broad Igf2 protein expression in littermate control wild type NEBs (+/+) and near-complete absence following *Igf2* deletion (–/–). n, NEBs scored; Cohen’s *d* −5.04, p<10^−4^. **(F)** Fraction of NE cells that proliferated in bronchial slice cultures (as in Fig. 1C-G) during treatment for 72 hours with recombinant Igf1 or Igf2 (100 ng/ml). Both ligands induced similar levels of NE proliferation above vehicle control (Cohen’s *d* 1.27-1.48, p<0.05). n, NE cells scored. **(G)** Scheme for testing autocrine Igf2 requirement for NE^stem^ proliferation after naphthalene injury. Tamoxifen induction of adult *Ascl1^CreERT2/+^; Igf2^flox/flox^; Rosa26^lsl-ZsGreen^* mice as in (B) deletes *Igf2* (and marks recombined cells with ZsGreen), and EdU injections track proliferation following naphthalene injury. **(H, I)** Representative photomicrographs of NEBs showing EdU incorporation (red) one week after injury in littermate control wild type NE cells (H, lacking *Ascl1^CreERT2/+^*) and conditional Igf2 deletion (I). Note reduction in EdU^pos^ NE cells in conditional *Igf2* deletion. Scale bars, 10 μm. **(J)** Quantification of (G-I) showing the fraction of NE cells per NEB that proliferated (EdU^pos^) by NE *Igf2* genotype. Conditional *Igf2* deletion reduced proliferation 76% (Cohen’s *d* −0.97, p<10^−7^). n, NEBs scored.

While Igf1 and Igf2 primarily engage Igf1r, they can also bind and function through the insulin receptor (Insr) and Igf1r-Insr heterodimers (*52*). *Insr* expression was detected (TPM>10) in 7-11% of NE cells (*17, 53*), mostly non-overlapping with *Igf1r-*expressors (68-71% *Insr^pos^* NE cells *Igf1r^neg^*, 88-96% *Igf1r^pos^* NE cells *Insr^neg^*) (fig. S4D). To determine if the residual NE^stem^ proliferation following airway injury is mediated by Insr, we conditionally deleted both *Igf1r* and *Insr* (*Insr^flox^* (*56*)) from NE cells (Fig. 2E). Although *Insr* deletion alone had little if any effect, deleting both receptors almost completely abolished NE^stem^ proliferation (mean 3% EdU^pos^ NE cells in compound homozygous mutants vs. 23.4% in controls, Cohen’s *d* −1.62, p<10^−5^ Mann-Whitney *U* test) (Fig. 2F-H). Thus, injury-induced proliferation of NE^stem^ is mediated by Igf1r and Insr.

### Igf2 is an autocrine mitogen for NE^stem^

Mouse lung expression patterns of *Igf1* and *Igf2* from scRNA-seq (*53*) are shown in Figure S3. Strikingly, *Igf2* is highly and selectively expressed in NE cells themselves (TPM>10 in 54% (*53*) and 69% (*17*) of NE cells), suggesting potential autocrine function. Immunostaining confirmed NE-selective localization of Igf2 protein (Fig. 3C, fig. S5A, B). All intrapulmonary NEBs and half of mini-clusters (*47*) expressed Igf2, and among clusters with at least one Igf2^pos^ cell, three-quarters of NE cells were Igf2^pos^ (NEBs 76%, mini-clusters 73%) (Fig. 3A). None of the neighboring epithelial cells, or the PSNs that innervate NEBs, or any other nearby cells in the lung showed reliable Igf2 labeling (fig. S5C-G), consistent with scRNA-seq. Solitary NE cells in the trachea, assessed by scRNA-seq (*57*) and by immunostaining, also did not express Igf2 (Fig. 3A, fig. S5H, I), nor did other tracheal cell types (fig. S5H, J-N). Clustered NE cells are thus the most prominent and perhaps exclusive source of Igf2 in mouse conducting airways (fig. S5O, P).

To test if Igf2 functions as an autocrine mitogen for NE^stem^ in airway repair, we conditionally deleted *Igf2* (*Igf2^flox^* (*58*)) from NE cells (Fig. 3B-E) and examined their proliferative response to injury as above (Fig. 3G). NE *Igf2* deletion reduced NE^stem^ proliferation by 76% (5% NE cells EdU^pos^ in *Igf2* homozygous mutants vs. 21% in wild type controls, Cohen’s *d* −0.97, p<10^−7^ Mann-Whitney *U* test) (Fig. 3H-J). A similar result obtained using the general epithelial driver *Shh^EGFP-Cre^* to delete *Igf2* in NE cells from the onset of epithelial development (63% reduction, Cohen’s *d* −1.17, p<10^−2^ Mann-Whitney *U* test) (fig. S7). These results parallel the effects of NE-specific *Igf1r* single and *Igf1r/Insr* double deletions (Fig. 2). We conclude that Igf2 functions as an autocrine mitogen in NE^stem^ during airway repair.

We also tested the effect of conditional deletion of *Igf1* (*Igf1^flox^*(*59*)) from two paracrine cell sources that express this alternative ligand: pulmonary mesenchyme (*Tbx4^LME-Cre^* (*60*)) (fig. S3A, notably adventitial fibroblasts), and pulmonary sensory neurons (*Phox2b^Cre^* (*61*)) (fig. S3B, notably PSN4 and PSN7 that directly innervate NEBs (*62*)). No significant effects on NE^stem^ proliferation were observed using the Cre drivers individually or together (fig. S8). Thus, paracrine Igf1 from these sources is dispensable for NE^stem^ proliferation following airway injury.

### Igfbp5 stabilizes and sequesters Igf2 in an inactive form within the niche

Expression patterns and gene deletion results described above show the mitogenic signal (Igf2), receptors (Igf1r, Insr), and signal transduction genes (e.g., Irs2 (*63*)) are all present in NE cells during homeostasis, and the pathway is fully functional as demonstrated by NE^stem^ proliferation following addition of Igf1 or Igf2 to lung slice cultures. This suggests that the activity of endogenously expressed Igf2 is normally somehow suppressed, since proliferation is nearly undetectable prior to injury. Circulating Igfs are typically bound by Igf binding proteins (Igfbps), a family of six proteins (Igfbp1-6) that bind Igfs with high affinity and, together in some cases with Igfals/ALS (*64*), stabilize the circulating hormones and keep them inactive until reaching their targets (*52*). We found that NE cells selectively express high levels of *Igfbp5* (75% NE cells TPM>10, all other *Igfbps* <4%) (fig. S3A), suggesting that a similar mechanism could operate locally within NEBs.

We first tested whether Igfbp5 stabilizes Igf2 in NEBs. In *Igfbp5* knockout mice, *Igf2* mRNA detected by smFISH in NE cells was unaltered (Fig. 4A, B). However, Igf2^pos^ NE cells in NEBs detectable by immunostaining were reduced by 75% (21% Igf2^pos^ NE cells in *Igfbp5* knockouts vs. 91% in wild type littermate controls, Cohen’s *d* −4.88, p<10^−6^ Mann-Whitney *U* test) (Fig. 4C-E). Thus, Igfbp5 stabilizes Igf2 protein in NEBs, presumably by forming a complex that prevents Igf2 degradation and diffusion out of the NEB.

**Figure 4.**
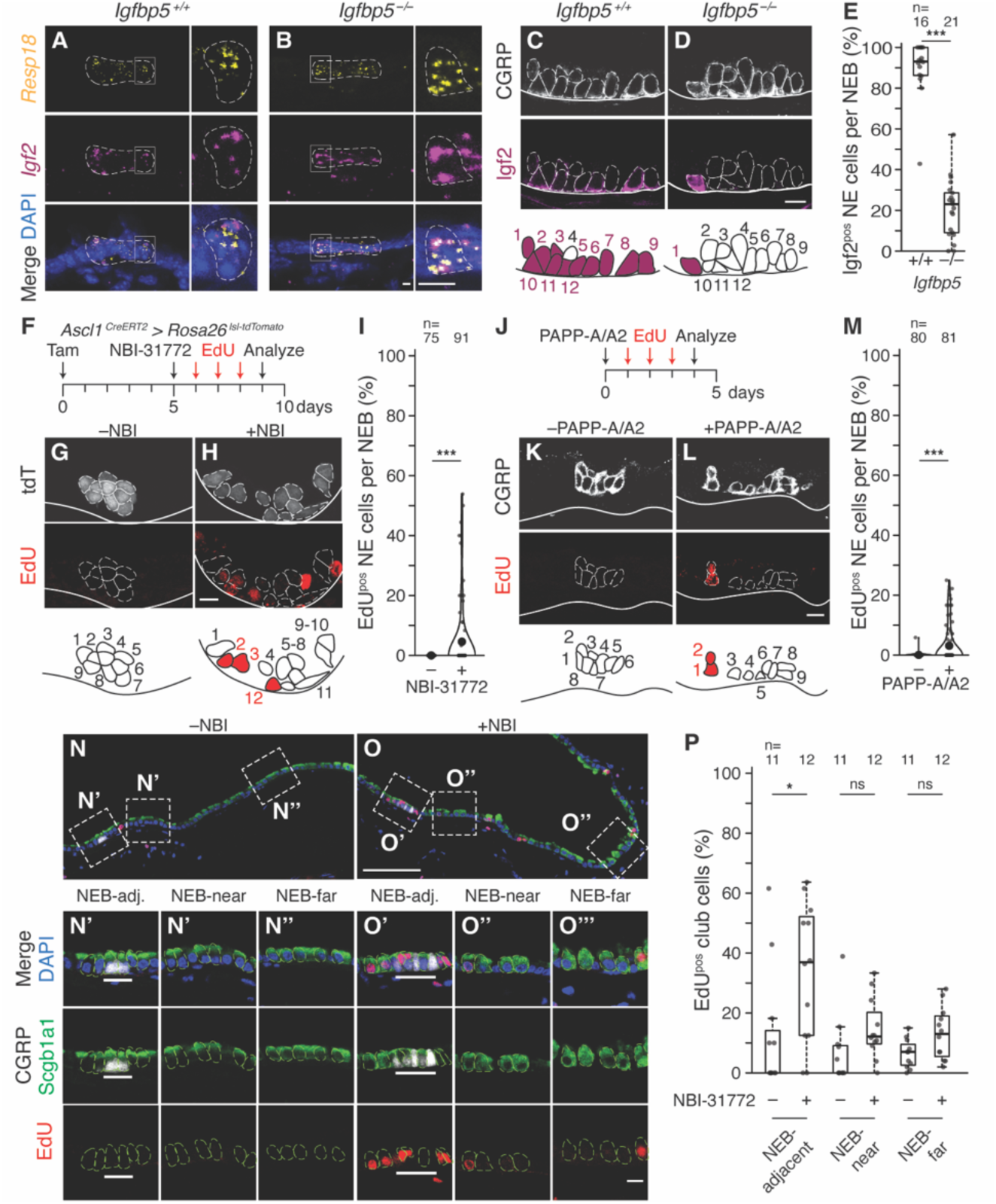
Effect of NE Igfbp deletion and pharmacologic disruption of Igf-Igfbp complexes on Igf2 protein stability and NE proliferation *in vivo*. **(A, B)** *Igf2* mRNA expression in NEB (left) and close-up (boxed area) showing individual NE cell (right) from adult *Igfbp5* wildtype littermate controls (*Igfbp5^+/+^*, A) and homozygous knockout mice (*Igfbp5^−/−^*, B), assessed by multiplex single molecule fluorescence in situ hybridization (smFISH, RNAscope v2) for *Igf2* (magenta) and NE marker (*123*) *Resp18* (yellow), with DAPI counterstain (blue). *Igf2* mRNA levels are not affected by *Igfbp5^−/−^*. Scale bars, 5 μm. **(C, D)** Photomicrographs and schematics of Igf2 protein in NEBs from adult *Igfbp5* wildtype (C) and homozygous knockout mice (D) assessed by immunostaining for Igf2 (magenta) and CGRP (white). Scale bars, 10 μm. **(E)** Quantification of (C, D) showing fraction of NE cells in each NEB with detected Igf2 by *Igfbp5* genotype. *Igfbp5* knockout reduced Igf2 staining by 75% (Cohen’s *d* −4.88, p<10^−6^). n, NEBs scored. **(F)** Scheme for testing effect on NE proliferation of acute, pharmacologic Igf/Igfbp complex dissociation. Tamoxifen induction of adult *Ascl1^CreERT2/+^; Rosa26^lsl-tdTomato/+^* mice labeled NE cells with tdTomato (tdT). Five days later the drug NBI-31772 (240 µg) was delivered by endotracheal instillation to acutely disrupt Igf/Igfbp complexes, then EdU was injected daily to track proliferation. **(G, H)** Photomicrographs and schematics of NEBs labeled with tdT (white) and EdU (red) four days after treatment with vehicle (G) or NBI-31772 (H). Scale bars, 10 μm. **(I)** Quantification of (F-H) as violin plots showing the fraction of NE cells per NEB that proliferated in each condition. Large black dots, distribution means. Note induction of NE proliferation by NBI-31772 (Cohen’s *d* 0.52, p<10^−3^). n, NEBs scored. **(J)** Scheme for testing effect on NE proliferation of acute Igf/Igfbp complex disruption by Igfbp proteases. Recombinant proteases PAPP-A (5 µg) and PAPP-A2 (5 µg) were delivered to adult C57BL/6 wild type mice by endotracheal instillation, then EdU was injected daily to track proliferation. **(K, L)** Photomicrographs and schematics of NEBs immunostained for CGRP (white) and EdU (red) four days after vehicle (K) or PAPP-A/PAPP-A2 protease (L) treatment. Scale bars, 10 μm. **(M)** Quantification of (J-L). Violin plots showing the fraction of NE cells per NEB that proliferated in each condition. Note induction of NE proliferation by protease treatment (Cohen’s *d* 0.69, p<10^−5^). n, NEBs scored. **(N, O)** NBI-31772 treatment also induces proliferation of nearby club cells. Photomicrographs of airways from same mice as (F-I), immunostained for EdU (proliferation, red), CGRP (NE cells, white), and Scgb1a1 (club cells, green, individual cells outlined in light green), with DAPI counterstain (blue). Boxes, closeups below showing club cells adjacent to NEBs (“NEB-adj.”, within two cell diameters of NEB) (N’, O’), near NEBs (“NEB-near”, two to ten cell diameters from NEB) (N’’, O’’), and far from NEBs (“NEB-far”, >20 cell diameters from NEB) (N’’’, O’’’). White line (in N’ and O’), NEB. Note prominent club cell proliferation adjacent to NEB after NBI-31772 treatment (O’). Scale bars, 100 μm (N, O), 10 μm (closeups). **(P)** Quantification of (N, O) showing the fraction of EdU^pos^ club cells under each condition and distance from NEBs. n, number of regions analyzed, with each datapoint representing the value for tens to hundreds of club cells within a scored region. NBI-31772 treatment induced proliferation in club cells adjacent to NEBs (Cohen’s *d* 0.93, p<0.05). ns, not significant.

To determine if acute release of Igf2 sequestered in an Igf2-Igfbp complex is sufficient to activate the NE^stem^ mitogenic pathway, we tested the effect of NBI-31772, a small molecule drug that releases Igfs from these complexes (*65, 66*). NBI-31772 was instilled endotracheally into uninjured mice in which NE cells were fluorescently labeled, and proliferation was tracked over four consecutive days by EdU injections (Fig. 4F). NBI-31772 treatment induced NE cell proliferation (4.6% NE cells EdU^pos^ vs. 0% in vehicle controls, Cohen’s *d* 0.52, p<10^−3^ Mann-Whitney *U* test) (Fig. 4G-I). Similar results obtained when NE cell proliferation was assessed by Ki67 immunostaining (*67*) at the endpoint (fig. S9A-D). Dissociation of Igf-Igfbp complexes is therefore sufficient to induce ectopic NE^stem^ proliferation.

The main physiological mechanism for release of circulating Igfs from Igf-Igfbp complexes once reaching a hormonal target site is proteolytic cleavage of the Igfbp (*52*). Many of the known Igfbp proteases are expressed by cell types located in and around NEBs (fig. S3), including the highly specific proteases PAPP-A (pregnancy-associated plasma protein A, also called pappalysin-1) and PAPP-A2 (pappalysin-2) (*68*). We tested whether endotracheal instillation of recombinant human PAPP-A/A2 could induce NE proliferation, as above for NBI-31772 (Fig. 4J). PAPP-A/A2 proteases also evoked NE proliferation, as measured by EdU incorporation (3.0% NE cells EdU^pos^ vs. 0.2% in vehicle controls, Cohen’s *d* 0.57-0.69, p<10^−5^-10^−3^ Dunn’s tests) (Fig. 4K-M) and Ki67 staining (3.2% Ki67^pos^ vs. 0.5% in controls, Cohen’s *d* 0.57-0.69, p<10^−5^-10^−3^ Dunn’s tests) (fig. S9E-H). Thus, proteolysis of Igfbps by PAPP-A/A2 proteases, like pharmacological dissociation of Igf-Igfbp complexes or airway injury, rapidly induces NE cell division.

In the Igf2-Igfbp complex disruption experiments, we also noted proliferation of rare airway epithelial cells surrounding NEBs (Fig. 4O, fig. S10). Their pattern suggested they might be a subpopulation of club cells identified previously to also function as stem cells after airway injury (*69, 70*). Immunostaining for Scgb1a1 (*71*) (also called CC10, CCSP, or uteroglobin) confirmed these as club cells, and showed their proliferation increased locally following drug treatment (37% of Scgb1a1^pos^ cells within two cell diameters of NEBs EdU^pos^ in NBI-31772 treated lungs vs. 14% in vehicle controls, Cohen’s *d* 0.93, p<0.03 Dunn’s test) (Fig. 4N-P). PAPP-A/A2 protease treatment gave a similar result, though the proliferative effect may have extended further (fig. S10). The results suggest that, in addition to its role as an autocrine mitogen acting on NE^stem^, released Igf2 may diffuse and influence proliferation of nearby club cells in a paracrine fashion, coordinating proliferation of NE^stem^ with other local stem cells after injury.

### Rb, but not p53, tonically prevents NE cell division downstream of Igf

Our original model (*17*) proposed that a hypothetical NE^stem^ mitogenic signal, identified here as Igf2, would act by transiently repressing the SCLC tumor suppressors Rb and p53, which together prevent NE^stem^ cell division under normal (non-injury) conditions. A prediction of this model is that ectopic activation of the “downstream” tumor suppressors (Rb and p53) should suppress proliferation of NE^stem^ induced by the “upstream” mitogen. We tested this molecular epistasis by supplying Igf1 or Igf2 to NEBs in lung slice cultures along with small molecule inhibitors Palbociclib, to inactivate Cyclin D-CDK4/6 and in turn activate Rb (*72*), and Nutlin-3a, to block binding of p53 to MDM2 and activate p53 (*73*) (Fig. 5A, B). Pharmacological activation in this way of either Rb or p53 (or both together) suppressed NE proliferation induced by Igf1 or Igf2 (Fig. 5C-L), indicating that Rb or p53, when active, are each able to block the mitogenic effect of Igfs. These tumor suppressors therefore function downstream or parallel to the Igf signaling pathway in NE^stem^.

**Figure 5.**
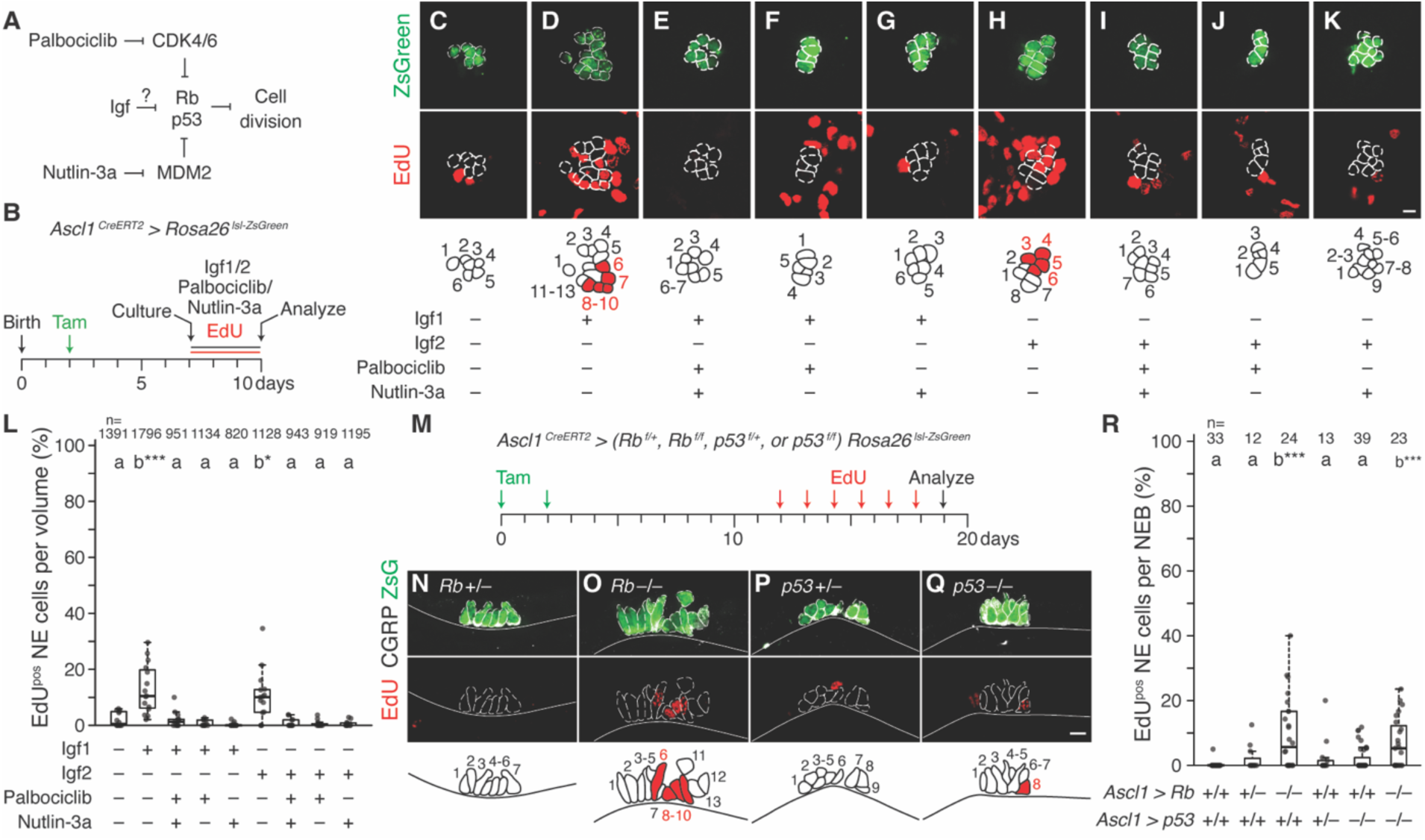
Genetic epistasis analysis of Igfs, Rb and p53 regulation of NE proliferation. **(A)** Genetic model: Igf pathway represses Rb and p53, which in turn repress NE cell division. Epistasis test: Model predicts Rb and p53 pharmacologic activation by Palbociclib and Nutlin-3a, respectively, should prevent cell division, even when Igf pathway is active. **(B)** Scheme for epistasis test in lung slice culture. Tamoxifen treatment of *Ascl1^CreERT2/+^; Rosa26^lsl-ZsGreen^* labels NE cells (ZsGreen), then P7 lung slice cultures are initiated with culture media supplemented throughout the full culture period with EdU, and with Igf1 (100 ng/ml), Igf2 (100 ng/ml), and tumor suppressor activators (Palbociclib 1μM, Nutlin-3a 10 μM) as indicated in panels (C-K). **(C-K)** Maximum intensity projections and schematics of representative NEBs stained for ZsGreen (NE marker) and EdU (proliferation marker) in lung slice culture epistasis test with culture medium supplemented as indicated. Note NE proliferation (EdU^pos^) in Igf1 and Igf2 treatment groups (D, H) but no proliferation in vehicle control (C) and all other conditions, including with Igf1 or Igf2 added but Rb and/or p53 activated by added Palbociclib and/or Nutlin-3a, respectively (E-G, I-K). Scale bars, 10 μm. **(L)** Quantification of (A-K) showing the fraction of NE cells that proliferated during each treatment. Statistical groupings determined by two-sided Dunn’s tests with Benjamini-Hochberg adjustments; asterisks denote significance relative to vehicle control (first). Note Rb or p53 when activated block Igf-induced NE cell proliferation. n, number of total NE cells scored. **(M)** Scheme for testing requirements of Rb and p53 to prevent NE cell proliferation under normal (uninjured) conditions. Tamoxifen induction of (N) *Ascl1^CreERT2/+^; Rb^flox/+^; Rosa26^lsl-ZsGreen/+^*, (O) *Ascl1^CreERT2/+^; R^bflox/flox^*; *Rosa26^lsl-ZsGreen/+^*, (P) *Ascl1^CreERT2/+^*; *p53^flox/+^; Rosa26^lsl-ZsGreen/+^*, or (Q) *Ascl1^CreERT2/+^; p53^flox/flox^; Rosa26^lsl-ZsGreen/+^* adult mice to excise essential exons from *Rb^flox^* and *p53^flox^* and label NE cells (ZsGreen). Eleven days later, daily EdU injections track proliferation over seven days. **(N-Q)** Photomicrographs and schematics showing NE cell proliferation (EdU^pos^) in representative NEBs (CGRP^pos^) of heterozygous or homozygous *Rb* (N, O) or *p53* (P, Q) NE conditional deletion mutants. Note increased proliferation only in homozygous *Rb* conditional deletion mutant (O). Scale bars, 10 μm. **(R)** Quantification of (M-Q) showing the fraction of NE cells per NEB that proliferated by NE *Rb* and *p53* NE conditional deletion genotype, including wild type (first) and *Rb/p53* compound mutant NEBs (last) assessed identically in Ref. (*17*) and shown here for comparison. Statistical groupings determined by two-sided Dunn’s tests with Benjamini-Hochberg adjustments; asterisks denote significance relative to wild type. n, NEBs scored. Note that *Rb* homozygous deletion in NE cells induced significant proliferation that matches the previously observed effect of *Rb/p53* compound deletion. By contrast, *p53* homozygous deletion alone induced little proliferation that was statistically grouped with wild type.

To determine if NE^stem^ require both Rb and p53 to maintain quiescence in uninjured, homeostatic conditions, as they do to prevent SCLC formation (*21, 74*), we deleted either *Rb* (*Rb1^flox^* (*75*)), or *p53* (*Trp53^flox^* (*76*)), or both together from NE cells (Fig. 5M, fig. S11A). NE *p53* deletion did not induce proliferation (Fig. 5P-R, fig. S11B, C, F). However, NE *Rb* deletion resulted in immediate cell division, and quantification showed that the amount of proliferation matched that of deleting both tumor suppressors together (*17*) (Fig. 5N, O, R, fig. S11D, F). Thus, Rb comprises the full proliferative check on NE^stem^ under homeostatic conditions.

Together with the above results using pharmacological activators of Rb and p53 in lung slice cultures, our findings demonstrate that Rb is tonically active during homeostasis to enforce NE^stem^ quiescence, and this cell cycle checkpoint functions downstream of and is inhibited by Igf signaling. By contrast p53, although also sufficient to block NE^stem^ proliferation when activated, remains inactive in NE^stem^ under homeostatic conditions. It appears to function as an NE^stem^ conditional checkpoint operating parallel to Igf signaling and the tonic Rb checkpoint (see Discussion).

### The two phases of the NE^stem^ cell repair program have distinct molecular controls

Immediately after injury, NE^stem^ proliferate for about a week, what we call the self-renewal phase (*17*). The repair program then transitions to a second phase in which a rare daughter cell (usually just one cell per NEB) undergoes reprogramming and transit amplification to generate a clone of highly proliferative epithelial daughter cells no longer of NE identity (*17*). These clonal outgrowths restore the injured epithelium surrounding NEBs by proliferating extensively for at least two more weeks, during which newborn cells eventually differentiate into appropriate mature airway epithelial cell types (*16, 17, 28, 29*) (e.g., club, ciliated, and goblet). The transition to this second phase is mediated by Notch (*17, 28*) and Yap (*29, 77*), likely (at least for Notch) activated by juxtacrine ligands expressed by adjacent surviving ciliated or club cells (*17*). To determine if these two phases of the repair program are contingently coupled, we assessed the second phase (clonal outgrowth formation) in *Igf1r/Insr* double deletion lungs that eliminate the first phase (self-renewal). We found that outgrowth frequency, assessed three weeks after injury (when outgrowths are large and easy to identify in whole mount lung preparations (*17, 28, 29*), Fig. 6A), was substantially (69%) reduced in the mutants (8.3% of NEBs formed outgrowths in mutants vs. 27.1% in controls, Cohen’s *d* −2.01, p<0.05 Mann-Whitney *U* test) (Fig. 6B-D). However, outgrowths that did form expanded a similar amount (Fig. 6 B, C, E). Thus, NE proliferation and self-renewal increases the likelihood that a clonal repair patch will form, but not the clonal growth rate and size of the repair patch.

**Figure 6.**
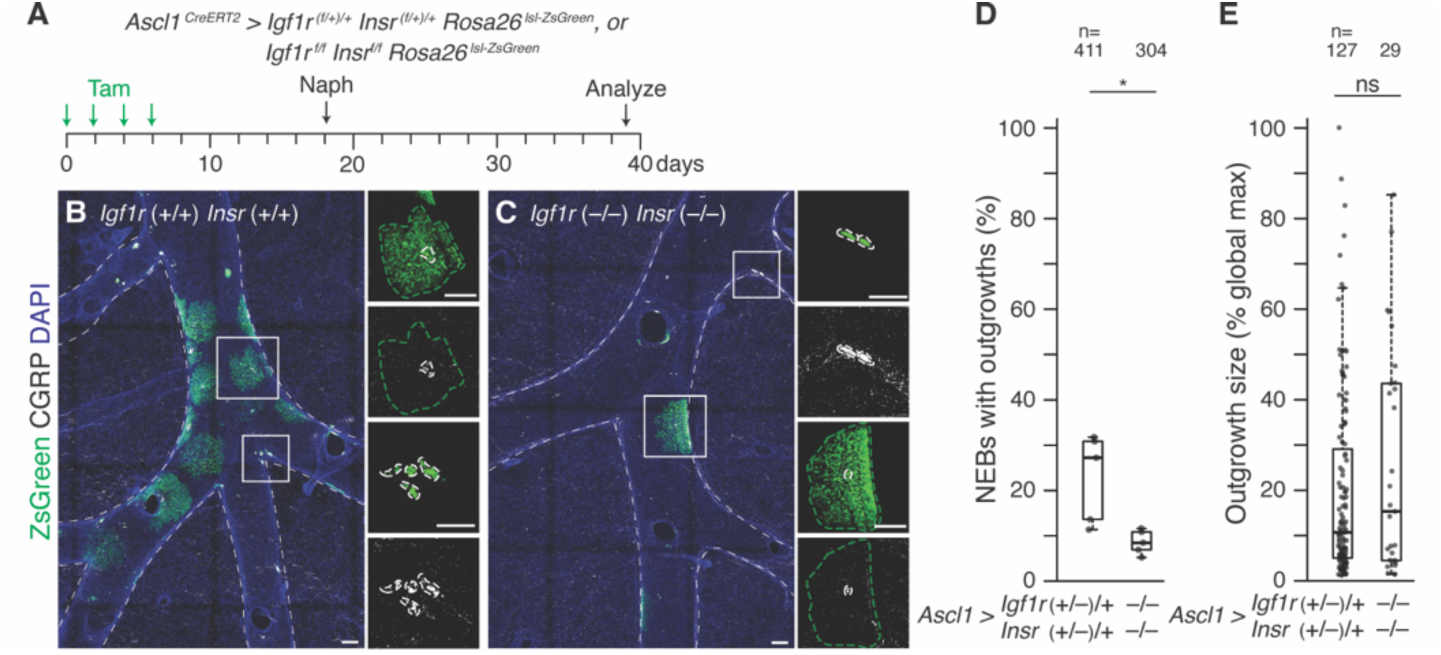
Distinct molecular controls the sequential self-renewal and reprogramming phases in the NE^stem^ sell repair program. **(A)** Scheme for testing *Igf1r* and *Insr* compound requirement for NE^stem^ reprogramming and outgrowth from NEBs, which begins about a week after naphthalene injury and forms large clonal epithelial repair patches surrounding NEBs (*17*). Tamoxifen induction of control (*Ascl1^CreERT2/+^; Igf1r^(flox/+)/+^; Insr^(flox/+)/+^; Rosa26^lsl-ZsGreen/ZsGreen^*) (B) or homozygous *Igf1r Insr* NE conditional deletion (*Ascl1^CreERT2/+^; Igf1r^flox/flox^; Igf1r^flox/flox^; Rosa26^lsl-ZsGreen/ZsGreen^*) adult mice (C). After 12 days to allow Cre recombination and Igf1r protein turnover, naphthalene was administered, and NEB outgrowths that form bronchial repair patches (defined as CGRP^neg^/ZsGreen^pos^ patches of epithelial cells surrounding NEBs) were scored three weeks later when outgrowths are large and mature. **(B-C)** Representative maximum intensity projections of bronchial branch z-stacks showing NEBs identified by CGRP immunostaining (white) and outgrowths by ZsGreen native fluorescence (green, green dashed outlines) in control (*Igf1r*+/+ *Insr+/+*) (B) and homozygous *Igf1r Insr* NE conditional deletion (*Igf1r*⎼/⎼ *Insr+/+*) (C) at three weeks after injury. DAPI (blue), counterstain for cell nucleus. Insets, close-ups of NEBs (boxed) without outgrowths (ZsGreen^pos^/CGRP^neg^, white dashed outlines) and with outgrowths (ZsGreen^pos^/CGRP^neg^, green dashed outlines) at three weeks. Scale bars, 100 μm. **(D)** Quantification of NEB expansion frequency three weeks after injury. Each dot represents counts from the right cranial lobe of an individual mouse. Conditional *Igf1r Insr* compound deletion reduces 69% of NEB expansion (Cohen’s *d* −2.01, p<0.05). n, NEBs scored. **(E)** Quantification of expansion outgrowth size three weeks after injury. Each black dot data point represents the size of a single outgrowth. Outgrowth size (in pixels) was normalized to the universal largest outgrowth identified. Note that both NEB outgrowth size at three weeks are similar between the two genotypes. n, number of expansion NEBs scored.

## Discussion

We identified the injury-activated mitogen and control pathway for NE^stem^ (Fig. 7A), stem cells that restore NEBs and large surrounding patches of bronchial epithelium following airway injury (*15–17*) and initiate small cell lung cancer (SCLC) following loss of Rb and p53 (*21, 74*). Surprisingly, the signal is autocrine Igf2, a classical circulating hormone in physiology (*52, 78*), growth (*79*), and aging (*80, 81*), but here is expressed by and acts locally on NE^stem^. The results support a model in which NE^stem^ serve as their own signaling niche (Fig. 7A, B), poised through selective and continuous expression of Igf2 and its receptors to generate an autocrine mitogenic signal, but constrained by the similarly selective and continuous expression of binding partner Igfbp5. Injury releases Igf2, perhaps by secretion of an Igfbp protease or protease activator by the injured cells. Igf2 thus freed activates its receptors on NE^stem^, triggering a signal transduction cascade that inactivates Rb, releasing this tonic NE^stem^ cell cycle checkpoint.

**Figure 7.**
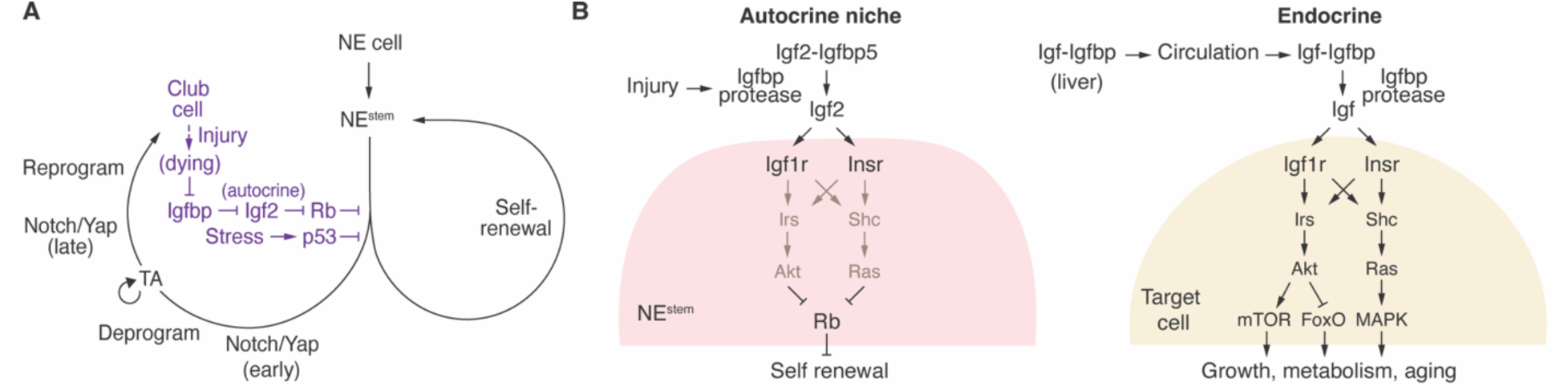
Genetic, cellular, and molecular models of NE^stem^ activation by injury. **(A)** Genetic logic. The multistep negative regulatory cascade poised for mitogenic activation of NE*^s^*^tem^ by airway injury, highlighting autocrine Igf2, the mitogen identified here with a central role in the cascade. Genetically upstream the mitogen is inhibited by autocrine Igfbp, but this inhibition is itself inhibited by airway (club cell) injury. Following disinhibition of Igf2 by injury, Igf2 autocrine signaling inhibits Rb, which itself normally inhibits NE^stem^ proliferation, maintaining their quiescence. Because cascade components are made constitutively, this double disinhibitory cascade is poised for rapid activation of NE^stem^ following injury. Dysregulation of other inhibitory steps such as Igfbp destruction or *Rb* deletion also rapidly activate NE^stem^, the latter we propose as the key initiating step in SCLC. p53 is another potent inhibitor of NE^stem^ activation that must be removed in progression to fulminant SCLC, but it is not normally operative in NE^stem^ and functions in parallel to Igf/Rb pathway, presumably activated by another cellular stress pathway that becomes active during the indolent phase as NE^stem^ slowly but continuously proliferates following loss of Rb. The Igf pathway is not required for the later steps of the stem cell program (e.g., deprogramming, reprogramming) that form NEB outgrowths and large clonal repair patches surrounding NEBs. Figure modified from Ref. (*17*) with newly identified features in purple. **(B)** Cellular and molecular model for autocrine Igf2 regulation of NE^stem^ in its niche (left) and comparison to classical Igf endocrine function (right). Autocrine niche function: During homeostasis, NE^stem^ produce Igf2 constitutively but it is stabilized in the niche and sequestered in an inactive (latent) complex with Igfbp, also produced constitutively. Airway injury triggers Igf2 release, perhaps by Igfbp destruction by PAPP-A/A2 proteases. Igf2 then engages and activates its receptors Igf1r and Insr, and this autocrine signaling leads to inhibition of Rb, releasing its inhibition of NE^stem^ self-renewal. Many of the same components are used as in the classical Igf endocrine pathway, but this autocrine niche pathway operates locally, is regulated by tissue injury, and the key regulatory event is disruption of the latent Igf-Igfbp complex and release of Igf. Endocrine function: Igf produced in the liver forms complexes with Igfbps that are secreted into circulation and distributed systemically before activation by Igfbp proteases at target tissues. Igf2 receptors (Igf1r, Insr) trigger downstream signaling pathways (e.g., Akt, Ras), which promote growth, metabolic processes, and aging. The system operates systemically, is generally regulated by physiological and developmental signals, and the key regulatory event is typically production or secretion of the Igf hormone at its source.

### Autocrine vs endocrine Igf signaling

This NE^stem^ control pathway has striking biochemical and molecular parallels to the classical Igf hormonal pathway (*52*) (Fig. 7B). The key distinguishing features are the source and regulation of the ligand. In the endocrine pathway, Igf ligand is produced far from its targets and stabilized by Igfbps as it circulates. The ligand is primarily regulated physiologically at the source via expression or secretion into circulation (*78, 81*). By contrast, in the NE^stem^ pathway, production of Igf is continuous and its actions local, within the niche and the stem cells it controls. The key regulatory event too is local, involving injury-induced release of Igf from the Igf-Igfbp complex. The regulation of this late step in the pathway is neatly tuned to its repair function, with the mitogenic ligand always present and poised for action immediately upon release from the complex by injury. Interestingly, the released Igf2 does not exclusively act cell autonomously. Nearby club cells also proliferate upon dissociation of the complex, suggesting that the Igf signal in the niche coordinates growth of both NE and surrounding stem cells (*69, 70*). Some of the released Igf2 could enter circulation and also have distant endocrine actions.

### Igf signaling and the initiation of small cell lung cancer

The results have important implications for SCLC. Prior studies of conditional *Rb* and *p53* double deletions in mouse lung show the prominent source of SCLC is NE cells (*74*), with NE^stem^ the most readily transformed into a continuously proliferative state (*17*). Our results show that Igf sigaling or *Rb* deletion alone is sufficient to activate NE^stem^ proliferation, creating slowly expanding clusters of NE cells. These do not progress to SCLC, however, because *Rb* deletion alone from NE cells does not generate fulminant SCLC tumors like those that arise when both *Rb* and *p53* are deleted (*82, 83*). We infer there is a subsequent NE^stem^ checkpoint mediated by p53, activated perhaps by genotoxic or metabolic stress encountered during the weeks of sustained proliferation following *Rb* deletion. According to this model, removal of both the p53-mediated conditional checkpoint and earlier Rb-mediated constitutive checkpoint are required for NE^stem^ clones to progress to malignant SCLC, an event that takes months in mice (*74*) and likely years or decades in humans (*21*). *Rb* homozygous mutant pediatric retinoblastomas also pass through an “indolence phase” of weeks to months (*84*), followed by a subsequent switch to aggressive malignancy that appears to also depend upon suppressing p53 (*85, 86*).

Expression and mutation signatures in human SCLC implicate Igf as an oncogenic driver: *IGF2* mRNA expression is high in tumor cells of some patients (*87*), a genomic locus including signal transducer *IRS2* exhibits focal and recurrent copy number gains (*30*), and mutations cluster in genes comprising the downstream PI3K pathway (*63, 88–90*) (e.g., *PI3KCA*, *PTEN*), of which *PTEN* is an especially potent SCLC tumor suppressor in mice (*91–93*). Although our NE^stem^ model predicts IGF signaling would be superfluous once downstream target *RB1* is deleted, upstream mutations could arise early (e.g., chronic cigarette smoke) and promote pre-cancerous growth (*94, 95*). IGF signaling might also contribute to the observed neural support of SCLC tumorigenesis (*96, 97*). Likewise, *IGF1* upregulation (*98*) could contribute to growth of carcinoids and other pulmonary NE neoplasms that resemble continuously cycling NE^stem^.

### Autocrine Igf signaling in diverse neuroendocrine cells and their tumors

Igfs are classically known as hormonal regulators of organismal growth, physiology, and longevity (*78, 79, 81*). Of the various cellular effects of Igfs that underlie these phenomena, proliferation was the first discovered and has been demonstrated for a variety of mammalian progenitors and cell types in culture (*52*). Although evidence for Igf function in adult stem cells is sparse, it has been implicated in neural stem cells in the hippocampus and subventricular zone (*99*) and some adult *Drosophila* stem cells (*100*). We suspect neurogenesis and production of NE cells is an ancient and general Igf function, shared across organs, organisms, and life stages, much like Egf signaling is broadly implicated in proliferation of epithelial progenitors (*35–43*). This idea fits nicely for pancreatic beta cells, which like pulmonary NE cells employ an autocrine Igf2-Igf1r-Irs2 signaling module to control their proliferation (*58, 101–105*), though activated by diet, pregnancy, and old age rather than injury.

The results suggest a special connection between Igf1r signaling and Rb. Although Rb is a vital cell cycle checkpoint and is sensitive to multiple growth factors (*106*), *RB1* mutations are only prevalent in a restricted set of cancer types including retinoblastoma (*107–109*), osteosarcoma (*107, 108*), small cell lung cancer (*110*), and a variety of other NE tumors (*111–113*). Similarly in mice, *Rb* deletion commonly induces tumors with NE features (*83, 114*), and Rb removal underlies the switch to a NE phenotype that occurs in some lung and prostate tumors of patients that gain chemotherapy resistance (*115, 116*), including to Egfr-targeted therapeutics in lung adenocarcinoma (*117–122*). These findings reveal a pronounced vulnerability in NE cells and their transformed progeny to cell cycle dysregulation caused by Rb dysfunction, which could be explained if proliferative control of NE cells by Igf2-Igf1r-Irs2 is mediated exclusively through Rb and not other cell cycle checkpoints. Detailed biochemical investigations in NE cells of the signaling cascade that couples Igf1r activation to Rb inactivation, the identities of relevant E2F partners, and the full battery of transcriptional switches induced upon Rb removal are now needed so that durable therapeutic strategies aimed at blocking Igf signaling and/or restoring Rb function in SCLC and other NE tumors can be achieved.

## Acknowledgments

We thank Tushar Desai, Jane Johnson, Julien Sage, and Bernard Thorens (mouse lines); Peng Li and Ria Sircar (reagents); Camille Ezran and Kyle Travaglini (help accessing and analyzing scRNA-seq data); Maria Petersen (figure design); Astrid Gillich, Christin Kuo, Ross Metzger, Julien Sage, and Laura Seeholzer (comments on manuscript); members of the Krasnow lab (experiments, reagents, interpretations, comments on manuscript). Mark A. Krasnow is a Howard Hughes Medical Institute Investigator. Requests for materials should be sent to Mark A. Krasnow.

## Funding

National Institutes of Health grant U01-HL099995

National Institutes of Health grant U01-CA23185

Howard Hughes Medical Institute

Virginia and D.K. Ludwig Fund for Cancer Research

National Science Foundation Graduate Research Fellowship

Stanford Graduate Fellowship

Stanford Cell and Molecular Biology Training Grant T32GM007276

## Author Contributions

Conceptualization: YZ, YO, MAK

Methodology: YZ, YO,

Visualization: YZ, YO

Investigation: YZ, YO, MMM

Writing: YZ, YO, MAK

Funding acquisition: YZ, YO, MAK

Unpublished Data: YL, MEK

Supervision: YO, MAK

## Data and materials availability

All reagents used in this study are available from the Lead Contact, Mark A. Krasnow, and will be provided upon request. All scRNA-seq datasets were published previously or have been posted to the open access preprint repository bioRxiv (RRID:SCR_003933), apart from mouse airway smooth muscle cells queried for *Igf2* expression. This paper does not report original code. Please see details in Methods.

