## supplements for "Stem cell control and cancer initiation by an autocrine, injury-activated Igf complex"

**The PDF file includes:**

Materials and Methods

Figs. S1 to S11

Tables S1 to S2

References (124-141)

### Materials and Methods

#### Animals

Mouse lines used were constitutive or tamoxifen-inducible Cre recombinase drivers *Ascl1*$^{CreERT2}$ (*50*) (RRID:IMSR_JAX:012882), *Shh*$^{EGFP\text{-}Cre}$ (*44*) (IMSR_JAX:005622), *Tbx4*$^{LME\text{-}Cre}$ (*60*) (IMSR_JAX:033331), *Phox2b*$^{Cre}$ (*61*) (IMSR_JAX:016223), and *Acta2*$^{CreERT2}$ (*124*) (IMSR_JAX:032758), Cre-dependent fluorescent reporters *Rosa26*$^{lsl\text{-}ZsGreen}$ (*51*) (Ai6, IMSR_JAX:007906) and *Rosa26*$^{lsl\text{-}tdTomato}$ (*51*) (Ai9, IMSR_JAX:007909), and Cre-dependent loss-of-function ("flox") alleles for *Egfr* (*45*) (MMRRC_031765-UNC), *Igf1r* (*55*) (IMSR_JAX:012251), *Insr* (*56*) (IMSR_JAX:006955), *Igf1* (*59*) (IMSR_JAX:012663), *Igf2* (*58*) (IMSR_JAX:032493), *Rb1* (*75*) (IMSR_JAX:008186), and *Trp53* (*76*) (IMSR_JAX:008462). *Igfbp5* knockout mice (strain T013950, RRID:SCR_017239) were obtained from GemPharmatech. Wild type inbred mice (6-10 weeks) were C57BL/6NCrl obtained from Charles River (strain 027, RRID:MGI:2683688). Genotyping was performed on tail clips utilizing oligonucleotide primers reported previously for each strain or provided by the vendor for *Igfbp5*. Adult mice aged 2-4 months were used in experiments (except early postnatal lung slice cultures, see below), with similar numbers of male and female mice analyzed and allocated evenly to experimental groups. Igf binding protein protease experiments (see below) utilized exclusively wild type male mice. Sample sizes are given in Figure Legends and in table S2, with 2-6 mice typically analyzed for each experimental condition and similar numbers for controls. Mice were maintained in a 12hr light/dark cycle with food and water provided ad libitum. All animal husbandry, maintenance, and experiments were approved by the Stanford University Institutional Animal Care and Use Committee (APLAC 9780, 26676, 33690).

#### Tamoxifen induction of Cre recombination

Tamoxifen (MilliporeSigma T5648) stock solutions (2 or 20 mg/ml) were prepared by sonication in corn oil and stored at -20°C. Intragastric injections of 0.1 mg (50 µl of the 2 mg/ml tamoxifen stock solution) were administered once to pups at postnatal day 2 (P2) for slice culture at P7. Intraperitoneal (i.p.) injections of 4 mg (200 µl of the 20 mg/ml stock solution) were administered to adult mice once or repeated once daily for the periods indicated in experimental schemes in Figures.

#### Airway injury with naphthalene

Naphthalene solution (50 mg/ml) was prepared immediately before use by dissolving naphthalene (Acros Organics AC180902500) in corn oil by gentle rocking at room temperature for 30-60 minutes, then passed through a 0.2 mm filter (Nalgene 723-2520) to remove any undissolved solute and sterilize. A single dose of naphthalene solution (275 mg/kg body weight) was delivered to adult mice by i.p. injection at least ten days after the final tamoxifen injection to allow tamoxifen clearance. Naphthalene-treated mice typically lost ~20%-25% of body weight in the three days after injury, then partially recovered to ~15%-20% loss at one week. Minorities of mice were either refractory to naphthalene (i.e., did not lose weight, appeared alert and well groomed, and were behaviorally active after injury), or extremely susceptible (i.e., lost >40% of body weight), and were excluded from analyses.

**Endotracheal instillation of NBI-31772, PAPP-A, and PAPP-A2**

Mice were subjected to anesthesia using 3% isoflurane in an induction chamber for a minimum of 15 minutes, then positioned on a Rodent Tilting Workstand (Hallowell EMC) for intubation. Illumination was directed onto the neck, the mouse's mandible raised, and the tongue extended using a Q-tip. The vocal cord was visualized with a speculum, and an intubation tube (1" 20G disposable catheter, Fisher Scientific) was introduced into the trachea using a guide wire (Mouse Intubation Pack, Hallowell EMC). Following guide wire removal, 60 µl of sterile 4 µg/µl NBI-31772 in PBS (R&D Systems 5192, 240 µg/mouse total dose), 30 µl 0.167 µg/µl recombinant human PAPP-A (R&D Systems 2487-ZNF, 5 µg/mouse) and 30 µl 0.167 µg/µl recombinant human PAPP-A2 (R&D Systems 1668-ZN, 5 µg/mouse), or 60 µl appropriate control solutions (10% (v/v) DMSO in PBS for NBI-31772; 20 mM Tris, 4 mM $CaCl_2$, 60 mM NaCl, 0.02% (w/v) CHAPS detergent in PBS for PAPP-A/A2) were administered into the lungs during natural breathing. The intubation tube was then gently removed, and the mouse was monitored for 30 minutes to ensure normal breathing during recovery.

**Cell proliferation analysis by EdU labeling**

For i.p. injections, the synthetic deoxyribonucleoside analog EdU (Thermo Fisher A10044) was dissolved at 2 mg/ml in sterile PBS (pH 7.4) and stored at -20°C. Adult mice were administered 0.2 mg (100 µl) doses daily as indicated in Figures, and no toxicity was apparent. For slice culture, EdU was dissolved at 10 mM (~2.5 mg/ml) in sterile PBS, stored at -20°C, and diluted 1,000-fold in culture medium for a 10 µM (~2.5 ng/ml) working concentration. EdU was detected following tissue fixation and immunostaining (see below) using click chemistry to covalently attach Alexa Fluor 488, 555, or 647 azide (Thermo Fisher C10337, C10338, C10340 Click-iT EdU imaging kits, or Thermo Fisher C10637, C10638, C10640 Click-iT Plus EdU imaging kits) to EdU alkyne incorporated into DNA during S phase of the cell cycle. Click reactions were allowed to proceed for 30 minutes on cryosections (20 µm) or 3 hours in slices (200 µm) in the presence of $CuSO_4$ catalyzer at room temperature, which was determined empirically to be sufficient for labeling reagents to fully penetrate and label tissues.

**Single-cell mRNA-seq (scRNA-seq) gene expression analysis**

All scRNA-seq datasets were published previously or have been posted to the open access preprint repository bioRxiv (RRID:SCR_003933), apart from mouse airway smooth muscle cells queried for *Igf2* expression (fig. S6A, see below). Briefly, scRNA-seq profiles for 100 adult mouse pulmonary NE cells (*17*) (GEO GSE136580) were generated using the Fluidigm $C_1$ microfluidic system (*125*) for single-cell capture of FACS-purified *Ascl1*$^{CreERT2}$ (*17*) or *CGRP*$^{CreERT2}$ (*16*) labeled cells and cDNA library preparation with Smart-seq (*126*), followed by read alignment to the mouse transcriptome with kallisto (*127*) (RRID:SCR_016582). Profiles for 162 pulmonary sensory neurons (PSNs) of 10 different types (*62*), including 15 NEB-innervating neurons of 2 different types, were generated by manually picking single neurons from nodose/jugular ganglia of mice following retrograde labeling from the lung, with Smart-seq for cDNA library preparation in plate format and transcriptome read alignment with kallisto. Profiles for 6,315 adult mouse lung cells of 41 different types (Tabula Muris Senis (*53*), https://tabula-muris-senis.ds.czbiohub.org/), including 180 NE cells identified by marker expression or *Ascl1*$^{CreERT2}$ labeling, were generated using FACS and cDNA library preparation with Smart-seq2 (*128*) in plate format and genome read alignment with STAR (*129*) (RRID:SCR_015899). (Note that mouse NE cells present in Tabula Muris Senis are also extensively described and analyzed by Kuo, et al. (*123*)). Profiles for 7,193 FACS-purified EpCAM$^{pos}$ adult mouse tracheal epithelial cells of 7 different types (*57*) (GEO GSE103354), including 96 NE cells identified by marker expression, were generated using the Chromium droplet-based system (3' Library v2, 10x Genomics) and CellRanger software (*130*) (v1.0.1, RRID:SCR_023221, 10x Genomics) for single-cell capture, cDNA library preparation, transcriptome read alignment, and UMI collapsing. Profiles for 1,962 adult mouse lung cells analyzed in fig. S6A, including 88 airway smooth muscle (ASM) cells identified by *Acta2*$^{CreERT2}$ labeling (*131*), were generated by FACS negative selection for Pecam1 (CD31) and Ptprc (CD45) followed by cDNA library preparation in plate format with Smart-seq2 and genome read alignment with STAR (Maya Kumar, personal communication). For Smart-seq and Smart-seq2 datasets, a sequencing depth-normalized expression value (either transcripts for kallisto alignments or counts for STAR alignments) of 10 per million mapped reads was used as the threshold to declare a gene "expressed" in a single cell, and cells with lower values were excluded from mean and breadth calculations used for dot plots. For 10x Genomics datasets the thresholds were 0.1 counts per ten thousand mapped reads or 0 UMIs.

**Identification of candidate mitogenic receptors in NE cells and their cognate ligands**

5,455 genes encoding predicted transmembrane proteins were retrieved from The Human Protein Atlas (*132*) (RRID:SCR_006710, protein class: Predicted Membrane Proteins) on April 9, 2017, and of the corresponding 5,068 mouse orthologues obtained using BioMart (*133*) (SCR_002987), 4,832 mouse genes were matchable to our NE scRNA-seq gene expression table (*17*). The 3,056 of these (63.2%) that were expressed in at least one of the 100 NE cells (TPM>10) were cross-referenced with a list of 77 receptors or receptor subunits curated to potentially control cell proliferation based on literature review (table S1), yielding 40 receptors/receptor subunits for further investigation in slice culture (see below). At least one cognate ligand or agonist per receptor or receptor family was identified based on literature review and obtained commercially (ligands in recombinant, soluble form): human Bdnf (R&D Systems 248-BDB, 100 ng/ml), Carbachol (Sigma C4382, 100 nM), mouse Fgf1 (Abcam ab73132, 100 ng/ml), human Gdnf (PeproTech 450-10, 10 ng/ml), human Hbegf (R&D Systems 259-HE, 50 ng/ml), mouse Igf1 (R&D Systems 791-MG, 100 ng/ml), ), mouse Igf2 (R&D Systems 792-MG, 100 ng/ml), Met-enkephalin (Sigma M6638, 1 μM), mouse Midkine (PeproTech 315-25, 100 ng/ml), mouse beta-Ngf (R&D Systems 1156-NG, 100 ng/ml), human Pleiotrophin (PeproTech 450-15, 100 ng/ml), mouse R-spondin1 (R&D Systems 7150-RS, 1 μg/ml), Smoothened agonist (SAG, Enzo ALX-270-426, 100 nM), mouse Scf (R&D Systems 455-MC, 100 ng/ml), mouse Wnt3a (PeproTech 315-20, 100 ng/ml). See table S1 for receptors grouped by signaling family, receptor expression levels in NE cells, ligand/receptor cognate relationships, working concentrations tested, carrier solutions, NE proliferation rates induced relative to vehicle control, sample sizes, and references. Two additional ligands, porcine gastrin-releasing peptide (Grp, R&D Systems 1788, 100 nM) and porcine neuromedin B (Nmb, R&D Systems 1908, 100 nM), were included due to their implication in SCLC despite their cognate receptors (Grpr, Nmbr) not being detected in mouse NE cell expression profiles.

**Early postnatal lung slice culture screen of candidate mitogenic ligands**

*Ascl1*$^{CreERT2/+}$*; Rosa26*$^{lsl\text{-}ZsGreen/+}$ or *Ascl1*$^{CreERT2/+}$*; Rosa26*$^{lsl\text{-}tdTomato/+}$ pups were genotyped the day after birth (P1) and NE cell labeling was induced in Cre-positive animals using intragastric tamoxifen injection at P2. At P7, 2-4 pups with NE labeling were euthanized by $CO_2$ asphyxiation and lungs dissected immediately following perfusion with ~2 ml sterile PBS into the right cardiac ventricle and intratracheal inflation with ~1 ml of 1.5% (w/v) low melting point (LMP) agarose (Invitrogen 16520) in sterile PBS. Lungs were kept protected from light in cold, sterile PBS until sectioning. The left lobe and right cranial lobe were embedded in 1.5% LMP agarose and fresh coronal slices (200 μm) prepared with a Compresstome vibrating microtome (Precisionary VF-200). Slices were quickly checked for presence of NEBs using an upright fluorescence stereomicroscope (Leica Biosystems MZ16 FA) and slices with NEBs were transferred to 24-well tissue culture plates (Falcon 353047) pre-coated with 200 μl of solidified, undiluted growth factor reduced Matrigel (Corning 354230). Plates were patterned such that each well contained two slices, each from a different individual pup, with two replicate wells per ligand, resulting in each ligand being tested on at least four slices from two individual mice per round of the experiment (repeated three times for a total of four rounds to test all candidate ligands, with replicate vehicle control wells included in each round). After placement on top of Matrigel, slices were fully submerged in culture medium consisting of DMEM/F12 with GlutaMAX (Gibco 10565018) and 10% (v/v) fetal bovine serum (FBS, Gibco 10082147) as the base medium, with EdU (10 μM) and one of each of the candidate recombinant ligands or agonists at a concentration taken from literature review (table S1). EdU and ligands/agonists were added to the base medium at 1,000-fold dilution in all cases except for Ephrin-A1/B1 (see below), R-spondin1 (50-fold dilution), and Wnt3a (100-fold dilution) to achieve desired working concentrations. In vehicle controls, sterile PBS containing 0.1% (v/v) bovine serum albumin (BSA) replaced the ligand or agonist. Under these conditions, slices exhibited minimal cell death after 72 hours as measured by exclusion of a plasma membrane impermeable fluorescent dye (DEAD red, Molecular Probes L-7013). Experimental cultures were incubated at 37°C for 72 hours (21% $O_2$, 5% $CO_2$), with replacement of the medium with identical fresh medium once halfway through the time course at 36 hours. Following culture, slices were washed briefly with PBS then fixed with 4% (v/v) paraformaldehyde (PFA, prepared fresh by dilution of a 32% stock solution (Electron Microscopy Sciences 15714) in PBS) for one hour at 4°C, before proceeding with EdU detection and DAPI staining (Invitrogen D1306, 100 ng/ml). Slices were optically cleared using the CUBIC method (*134*), comprising a one-hour incubation in CUBIC 1 reagent at room temperature with gentle rocking and storage in CUBIC 2 reagent at 4°C until microscopy (see below). Native ZsGreen or tdTomato fluorescence was used to identify and score NE cells.

**Ephrin-A1 and Ephrin-B1 molecular clustering**

Recombinant Ephrin-A1 (R&D Systems 602-A1) and Ephrin-B1 (R&D Systems 473-EB) Fc chimera proteins (consisting of mouse Ephrin extracellular domains fused to human $IgG_1$ heavy chain) and donkey anti-human $IgG_1$ (Jackson ImmunoResearch 709-005-098) were reconstituted in sterile PBS to 100 μg/ml. 10 μl of each Ephrin Fc was incubated with 2 μl anti-$IgG_1$ (5:1 molar ratio) for 30 minutes at room temperature with rotation to induce 2:1 Ephrin Fc:anti-$IgG_1$ molecular clustering, as previously described (*135*) (each of the two anti-$IgG_1$ variable regions binds one Ephrin Fc). This yielded 83.3 μg/ml clustered Ephrin Fc stock solutions which were added to slice culture base medium at 83-fold dilution for a final working concentration of 1 μg/ml. 0.2 μg/ml anti-$IgG_1$ in PBS (the same concentration present in clustered solutions) was used as a negative control.

**Molecular epistasis between Igf ligands and Rb/p53 tumor suppressors**

Fresh lung slices (200 μm) were collected at P7 from *Ascl1*$^{CreERT2/+}$*; Rosa26*$^{lsl\text{-}tdTomato/+}$ pups that received tamoxifen at P2 and were prepared for culture on Matrigel as above. Culture medium consisted of DMEM/F12 with GlutaMAX, 10% FBS, and 10 μM EdU as the base medium, with recombinant mouse Igf1 (R&D Systems 791-MG, 100 ng/ml), recombinant mouse Igf2 (R&D Systems 792-MG, 100 ng/ml), Palbociclib (MilliporeSigma PZ0383, 1 μM), and Nutlin-3a (MilliporeSigma SML0580, 10 μM) applied by dilution of 1,000-fold concentrated stock solutions. DMSO (1,000-fold diluted) was the vehicle control. Following the 72-hour culture period (37°C, 21% $O_2$, 5% $CO_2$, with replacement with identical fresh medium once at 36 hours), slices were fixed, stained for EdU and DAPI, and optically cleared using CUBIC as above. Native tdTomato fluorescence was used to identify and score NE cells.

**Immunostaining**

Mice were euthanized by $CO_2$ asphyxiation and lungs (in some cases with the trachea purposely still attached) were dissected immediately following perfusion with ~5ml PBS into the right cardiac ventricle and intratracheal inflation with ~2ml of 2% LMP agarose in PBS. Lungs were fixed in 4% PFA for six hours at 4°C with gentle rocking and washed into cold PBS kept protected from light for short-term storage at 4°C.

For cryosections, lungs were dehydrated in 30% (w/v) sucrose in PBS overnight or until they sank at 4°C before embedding in optimum cutting temperature compound (OCT, Tissue-Tek 4583) and storage at -80°C. Lungs and trachea were embedded separately, with the dorsal surface facing down for lungs and with the long axis lying horizontally for tracheae. Coronal cryosections (20 μm) were prepared using a cryostat (Leica Biosystems CM3050 S) and were adhered to Superfrost Plus slides (VWR 48311-703), dried at room temperature for 15-60 minutes, and washed twice at room temperature with gentle rocking for five minutes each in 0.3% (v/v) Triton X-100 (Sigma X100) in PBS. Sections were incubated for one hour at room temperature in either 5% (v/v) normal goat (Jackson ImmunoResearch 005-000-121) or donkey serum (Jackson ImmunoResearch 017-000-121) in 0.3% Triton X-100 PBS (cryoblock solution), then incubated overnight at 4°C in cryoblock containing primary antibodies: mouse monoclonal anti-Acta2-Cy3 (AB_476856, 1:200), guinea pig polyclonal anti-CGRP (EuroProxima 2263B-GP470-1, diluted 1:1000), rabbit polyclonal anti-CGRP (RRID:AB_2068527, 1:1000), sheep polyclonal anti-CGRP (AB_2314159, 1:1000), rabbit polyclonal anti-Col1a1 (AB_2547045, 1:500), rat monoclonal anti-E-cadherin (AB_2533005, 1:500), goat polyclonal anti-Igf2 (AB_2122526, 1:100), goat polyclonal anti-Itga8 (AB_2296280, 1:500), rat monoclonal anti-Ki67-eFluor 660 (AB_2574235, 1:250), chicken polyclonal anti-Krt5 (AB_2565054, 1:500), rat monoclonal anti-Pecam1 (AB_394816, 1:500), rabbit polyclonal anti-Scgb1a1 (AB_310759, 1:500), and rabbit monoclonal anti-Tuj1 (AB_2566588, 1:500). The following day, sections were washed three times for five minutes each in 0.3% Triton X-100 PBS, then incubated for one hour at room temperature in cryoblock containing Alexa Fluor-conjugated secondary antibodies (Thermo Fisher goat or Jackson ImmunoResearch donkey anti-IgG polyclonals (anti-$IgG_1$ for mouse monoclonal primaries) conjugated to Alexa Fluor 488, 555, 568, 633, or 647, diluted 1:200-500) and DAPI (100 ng/ml). Stained sections were finally washed twice for five minutes each in 0.3% Triton X-100 PBS and once for five minutes in PBS, then mounted on coverslips (VWR 48393-106) using Mowiol 4-88 (MilliporeSigma 81381) with DABCO antifade (MilliporeSigma D2522, prepared according to http://cshprotocols.cshlp.org/content/2006/1/pdb.rec10255 except the centrifugation step) or Fluromount-G (SouthernBiotech 0100-01) as the mounting medium. Mounted specimens were stored at 4°C until microscopy.

For whole-mount preparations, coronal sections (500 μm) of PFA-fixed left lobes were prepared with a vibrating blade microtome (Leica Biosystems VT1000 S). Sections were incubated overnight at 4°C with gentle rocking in 5% normal goat serum in 0.5% Triton X-100 PBS (whole mount block solution), then incubated in whole mount block containing unconjugated rabbit polyclonal anti-CGRP (AB_2068527, 1:1000) for four days at 4°C with gentle rocking and light protection. Primary-stained slices were washed in 0.5% Triton X-100 PBS 5-6 times for one hour each at room temperature with gentle rocking, then incubated in whole mount block containing Alexa Fluor 555-conjugated goat polyclonal anti-rabbit IgG (AB_2535850, 1:500), and DAPI (100 ng/ml) for two days at 4°C with gentle rocking and light protection. Secondary-stained slices were washed again as above then optically cleared using overnight incubation in CUBIC-L (TGI T3740) at room temperature with gentle rocking and storage in CUBIC-R+(M) (TGI T3741) at 4°C until confocal microscopy.

***Igf1r* detection in NEBs by single molecule fluorescence in situ hybridization (smFISH)**

Mice were euthanized and lungs were dissected and prepared for fluorescent *in situ* hybridization with RNAscope (*136*) (Advanced Cell Diagnostics) as for immunostaining, except that nuclease-free buffers were used for all incubations and washes. Coronal cryosections (12 μm) were adhered to Superfrost Plus slides, dried at room temperature for 15-30 minutes, pretreated by incubating slides in Target Retrieval buffer (ACD 322000) at 95°C for five minutes, washed twice with nuclease-free water and once with 100% ethanol for two minutes each, then treated with Protease III (ACD 322281) for 30 minutes at 40°C. Slides were then washed twice with nuclease-free water before proceeding with mouse *Igf1r* probe hybridization (ACD 417561-C2, delivered in probe diluent) for two hours at 40°C and non-enzymatic amplification of puncta using the RNAscope Multiplex Fluorescent Reagent Kit v1 (ACD 320850). Following completion of the smFISH protocol, tdTomato expression resulting from tamoxifen induction of $Ascl1^{CreERT2}$ was detected by immunostaining using a rabbit polyclonal anti-RFP (AB_2209751, 1:500), then finally EdU by click chemistry (AF647) and cell nuclei with DAPI before mounting coverslips with Mowiol 4-88/DABCO and proceeding to confocal microscopy. smFISH puncta were scored only if signal was detected in multiple adjoining pixels in three dimensions (i.e, within an optical section or in consecutive optical sections) to avoid tallying spurious puncta possibly due to imperfect camera function.

#### *Igf2* smFISH in NEBs and airway smooth muscle cells

Lung coronal cryosections (10 μm) were adhered to Superfrost Plus slides and prepared for smFISH according to the manufacturer's instructions detailed in the RNAscope Multiplex Fluorescent Reagent Kit v2 (ACD 323100). Mouse *Igf2* probe (437671-C1) was combined with either *Resp18* (493871-C3) (a sensitive and specific marker for mouse NE cells (*123*)) for visualization in NEBs, or *Acta2* (319531-C2) in airway smooth muscle cells (*60*). ACD kit instructions were followed and the smFISH protocol completed (including probe hybridization, non-enzymatic signal amplification, and HRP signal development) before proceeding with immunostaining (CGRP), DAPI counterstaining, and imaging.

#### Microscopy and imaging

Cryosections were imaged using a Zeiss LSM 780 or Leica Stellaris 8 laser scanning confocal microscope with inverted 40X oil immersion objectives (Carl Zeiss AG, NA=1.4; Leica HC PL APO, NA=1.3). Optical sections were collected at 0.5 μm resolution through the z plane (dorsoventral axis of the lung). Images described in Figure Legends as "photomicrographs" are single optical sections, but image feature quantifications represent cumulative counts through multiple optical sections. Serial sections were generally visually inspected to ensure that no apparent differences in the various feature patterns reported were observed, however for consistency serial sections of the same NEB were excluded from quantifications. Except when explicitly comparing Igf2 expression in NEBs, mini-clusters, and tracheal singletons, NEBs scored ranged from 5 to 137 NE cells (median 16, mean 19.4), and therefore included the upper range of what are defined as mini-clusters (*47*). Scored NEBs were anatomically located within the pulmonary bronchi and bronchioles (termed "airways").

Cultured lung slices were placed in inverted coverglass chambers (Nunc Lab-Tek 155361), mounted with coverslips (VWR 89015-724) using CUBIC 2 as the mounting medium, and imaged using a Zeiss LSM 780 laser scanning confocal microscope with an inverted 25X oil immersion objective (Carl Zeiss AG, NA=0.8). Optical sections were collected at 2 μm resolution. $EdU^{pos}$ NE cells were scored in all NE cells present (including NEBs, mini-clusters, and singletons) in individually imaged volumes (567 μm x 567 μm x 200 μm), and the ligand-induced proliferation rate calculated by dividing $EdU^{pos}$ NE cells by the total number of NE cells present in the volume.

Whole mount lung sections were placed in inverted coverglass chambers, mounted with coverslips, and imaged using a Leica Stellaris 8 laser scanning confocal microscope with an inverted 10X air objective (Leica HCX PL, NA=0.30). Optical sections were collected at 6 µm resolution. NEB outgrowth sizes and frequencies were measured in whole left lobes from mice three weeks after injury. Images were intensity thresholded to remove background fluorescence and contiguous signals (representing NEBs and associated outgrowths) were segmented and measured in units of pixels for each NEB with or without expansion.

Fiji (*137*) (RRID:SCR_002285) was used to color, overlay, adjust brightness and contrast, and add scale bars to images. The Cell Counter software plugin was used to aid manual counts of features of interest. Adobe Illustrator 2024 (SCR_010279) was used to rotate, crop, and annotate images.

**Statistical analysis**

Sample sizes (n) including the number of NE cells, number of NEBs, and number of mice analyzed are reported for each experiment in Figures, Figure Legends, and in table S2. Boxplots (also called box-and-whisker plots) show distribution median in bold, interquartile range (IQR, box), and range within 1.5 times the IQR (whiskers), with all data points overlaid. Violin plots show mirrored density distributions across the full range and the mean value in bold, with all data points overlaid. Plots were generated and data analyzed in R version 4.2.2 (RRID:SCR_001905) and RStudio version 2023.03.1+446 (SCR_000432). Effect sizes between pairs of distributions were calculated using Cohen's *d*, implemented using the *cohen.d* function from the *effsize* package. Statistical significance between pairs of vector distributions was calculated using the nonparametric Mann-Whitney *U* test (also known as the Wilcoxon rank-sum test), implemented using the *wilcox.test* function from the base *stats* package. Multiple vector distributions were compared with nonparametric Kruskal-Wallis *H* tests (also known as one-way ANOVA on ranks), using the *kruskal.test* function (*stats*), followed by Dunn's test to explicitly compare distributions pairwise, using the function *dunnTest* (*FSA*). Binomial and Fisher's exact tests were implemented using the *binom.test* and *fisher.test* functions, respectively (*stats*). Multiple hypothesis adjustments were made by the Benjamini-Hochberg procedure, using the *p.adjust* function (*stats*). Asterisks are used to denote significance, with *$p<0.05$, **$p<0.01$, and ***$p<0.001$.

### Fig. S1

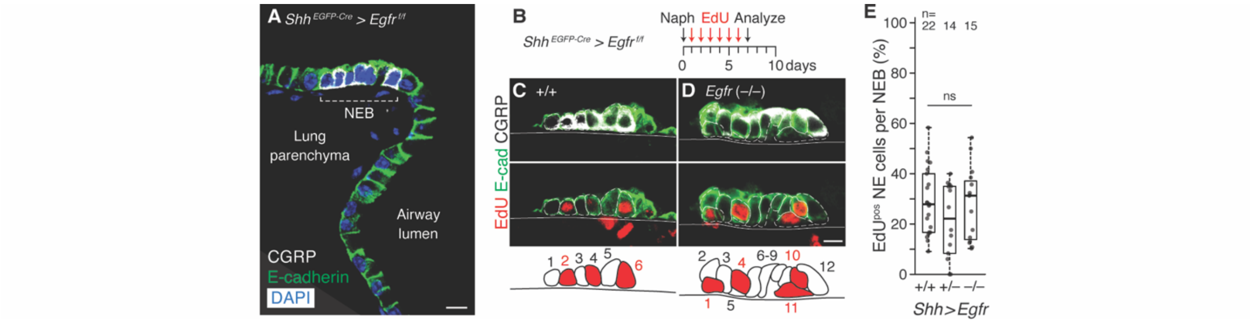


**Figure S1. Egfr is dispensable for $NE^{stem}$ proliferation after airway injury.**

**(A)** Photomicrograph of a neuroepithelial body (NEB) at a bronchial branchpoint in an adult *$Shh^{EGFP\text{-}Cre/+}$; $Egfr^{flox/flox}$* male mouse (36 weeks old) co-stained for CGRP (NE cells, white) and E-cadherin (airway epithelium, green) and counterstained with DAPI (cell nuclei, blue). *$Shh^{EGFP\text{-}Cre}$* excises an essential exon from both alleles of *$Egfr^{flox}$* in pulmonary epithelium (including NE cells) during embryogenesis (*138*). Conditional deletion NEB has normal NE cell number and morphology. Scale bar, 10 µm.

**(B)** Scheme for testing Egfr requirement for $NE^{stem}$ proliferation following airway injury with naphthalene. Experimental genotype as in (A) (*$Egfr^{flox/flox}$*, denoted –/–) along with *Egfr* wild type (*$Egfr^{+/+}$*, +/+) and heterozygous (*$Egfr^{flox/+}$*, +/–) littermate controls. Daily EdU injections track cumulative proliferation for one week after naphthalene (Naph) injury of airway.

**(C, D)** Photomicrographs and schematics of NEBs of wild type (C) and homozygous epithelial *Egfr* deletion (D) immunostained one week after injury for CGRP (white), E-cadherin (green), and EdU (red). Top row, CGRP/E-cadherin merge; bottom row, E-cadherin/EdU merge. Note *Egfr* deletion did not reduce NE cell proliferation. Scale bars, 10 µm.

**(E)** Quantification of (B-D) showing the fraction of NE cells per NEB that proliferated by epithelial *Egfr* genotype as boxplots (median values shown as thick horizontal lines). ns, not significant (see table S2 for statistical test applied and values obtained in all plots).

**Fig. S2**

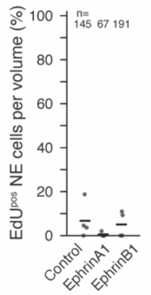


**Figure S2. Testing mitogenic effect on NE cells of EphrinA1 and B1 in postnatal lung slice culture.** Experimental animals, procedure, and analysis identical to Fig. 1C-E. Neither Ephrin had a significant mitogenic effect.

### Fig. S3

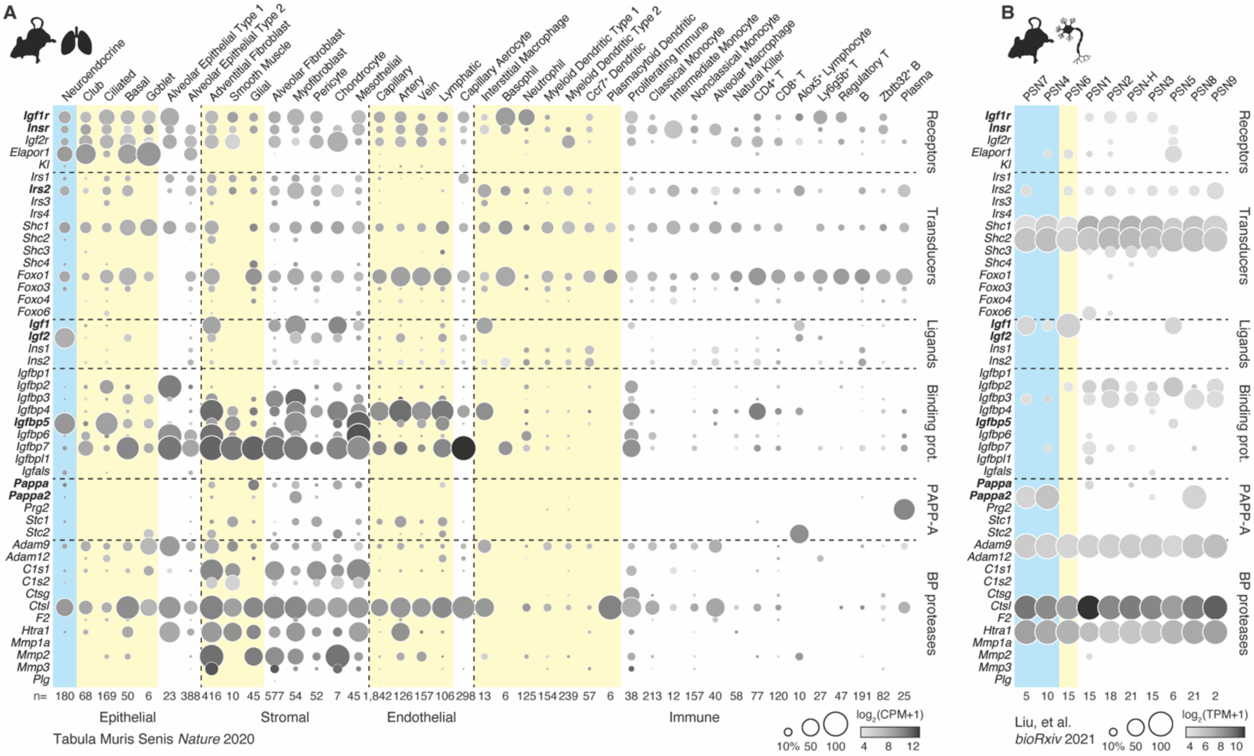


**Figure S3. Gene expression patterns of Igf signaling pathway components in adult mouse lung and pulmonary sensory neurons (PSNs) assessed by scRNA-seq.**

Mean level and breadth of expression of Igf signaling pathway components in adult mouse lung (*53*) (A, Smart-seq2) and pulmonary sensory neurons (*62*) (PSNs) (B, Smart-seq). Genes indicated at left are arranged according to protein classes (top to bottom: receptors, transducers, binding proteins, PAPP-A proteases and inhibitors, binding protein (BP) proteases) separated by horizontal dashed lines and ordered within each class alphanumerically, with genes of special interest highlighted in bold. Cell types are columns arranged in (A) according to tissue compartment (epithelial, stromal, endothelial, immune) separated by vertical dashed lines, then ordered within each compartment so that NE cells appear first (cyan shading) and other cell types predicted to be nearby NEBs next (yellow shading). For PSNs (B), neuron types that directly innervate NEBs (PSN7, PSN4) are listed first (cyan shading), and PSN6 which innervates smooth muscle preferentially at bronchial branch points where NEBs are enriched, next (yellow shading). Remaining PSNs are arranged alphanumerically and are either known to terminate distant to NEBs or have uncharacterized innervation patterns in the lung. PSN-H ("hybrid") has expression features in common with both PSN2 and PSN3. Number of cells per type is shown below each column. Dot intensity, expression level; dot size, breadth of expression in cell type; CPM, counts per million mapped reads; TPM, transcripts per million mapped reads. Note that dot intensity represents different methods of expression quantification and expression levels in the two plots (A and B).

**Fig. S4**

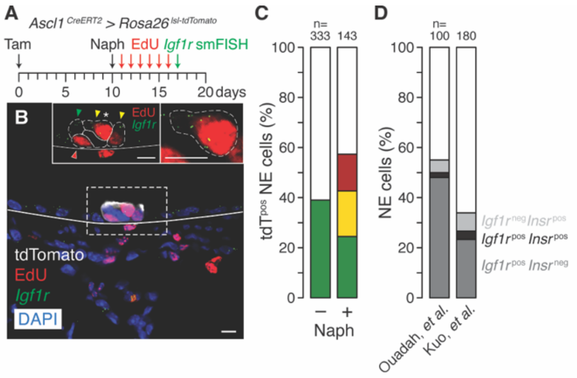


**Figure S4. Igf1 receptor marks proliferative NE cells after airway injury.**

**(A)** Scheme for assessing proliferation of *Igf1r*$^{pos}$ NE cells following airway injury with naphthalene. Tamoxifen (Tam) induction of *Ascl1*$^{CreERT2/+}$*; Rosa26*$^{lsl\text{-}tdTomato}$ adult mice labels NE cells (*17*) with tdTomato ten days prior to naphthalene (Naph) injury. One week after injury, NEBs are analyzed for *Igf1r* mRNA expression by single molecule fluorescence *in situ* hybridization (smFISH, RNAscope v1). Daily EdU injections track cumulative proliferation for the repair duration.

**(B)** Photomicrograph of a NEB as above 14 days after injury and probed for *Igf1r* RNA (green dots) costained for tdTomato (NE cells, white) and EdU (red) and counterstained with DAPI (blue). Of the five NE cells shown, three are EdU$^{pos}$ (red nuclei indicated by red and yellow arrowheads), three are *Igf1r*$^{pos}$ and partially overlap with EdU$^{pos}$ (green puncta indicated by green and yellow arrowheads), and one is EdU$^{neg}$/*Igf1r*$^{neg}$ (no arrowhead). Left inset, close-up of NEB showing EdU and *Igf1r* signals with individual NE cells outlined. Right inset, close-up of a single EdU$^{pos}$/*Igf1r*$^{pos}$ NE cell (denoted by asterisk in left inset). Scale bars, 10 μm.

**(C)** Quantification of (A, B) showing colocalization distributions for EdU incorporation and *Igf1r* expression in NE cells from NEBs before (–Naph) and one week after (+Naph) naphthalene injury. White, EdU$^{neg}$/*Igf1r*$^{neg}$ (61% before, 43% after); red, EdU$^{pos}$/*Igf1r*$^{neg}$ (0%, 15%); yellow, EdU$^{pos}$/*Igf1r*$^{pos}$ (0%, 18%); green, EdU$^{neg}$/*Igf1r*$^{pos}$ (39%, 24%). EdU$^{pos}$ cells are 1.7-times more likely to express *Igf1r* than not (42.6% vs. 25.6%, $p<0.05$ Fisher's exact test). The EdU$^{pos}$/*Igf1r*$^{neg}$ are presumably daughter cells that turned off *Igf1r* expression following proliferation of the mother cell.

**(D)** Quantification *Igf1r* and *Insr* mRNA expression in adult mouse NE cells from our two scRNA-seq datasets (*17, 123*) (280 total cells). White, *Igf1r*$^{neg}$/*Insr*$^{neg}$ (45%, 66%); light gray, *Igf1r*$^{neg}$/*Insr*$^{pos}$ (5%, 7%); black, *Igf1r*$^{pos}$/*Insr*$^{pos}$ (2%, 3%); dark gray, *Igf1r*$^{pos}$/*Insr*$^{neg}$ (48%, 24%). Note rare *Igf1r/Insr* expression overlap in NE cells.

### Fig. S5

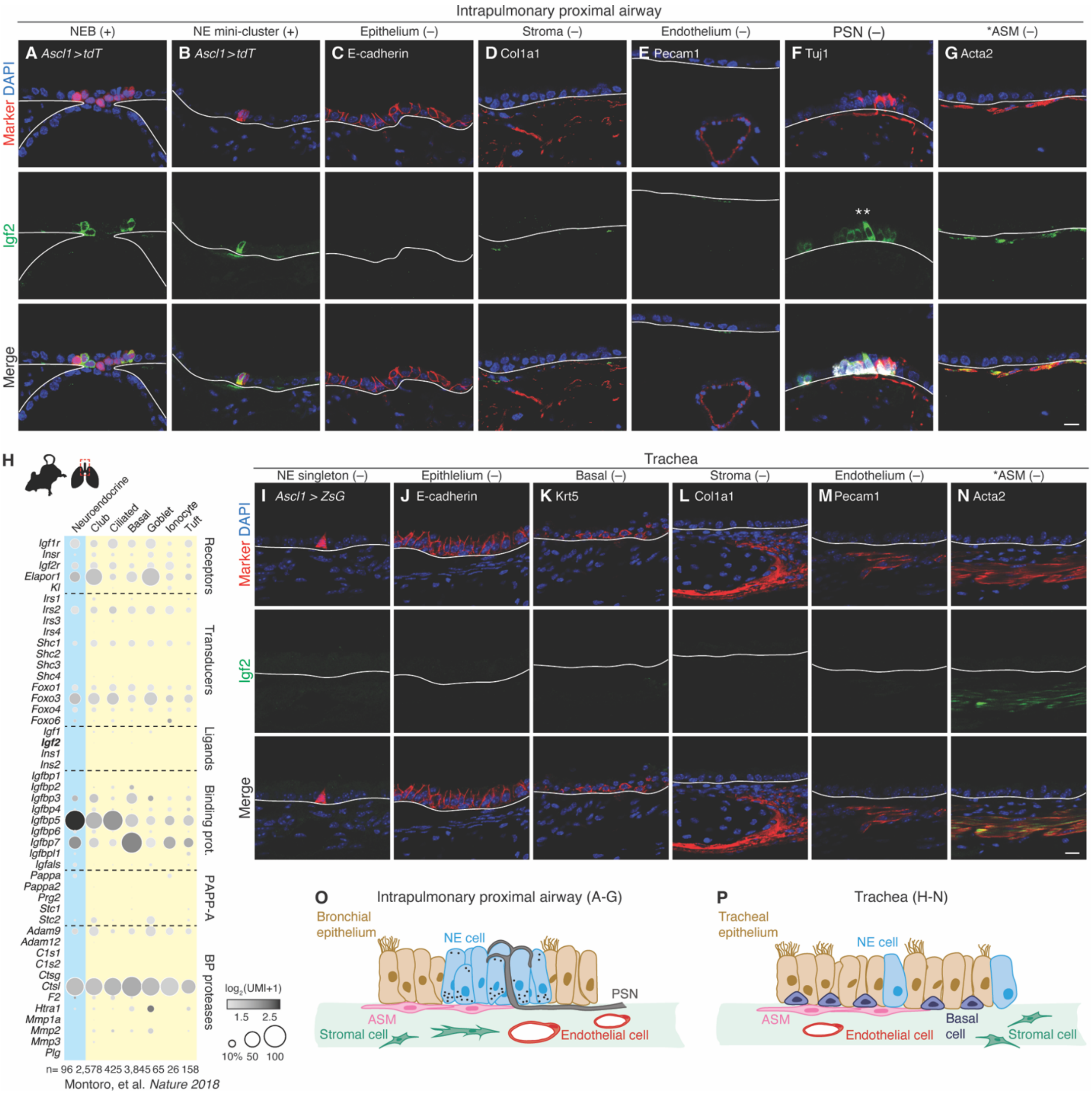


**Figure S5. Igf2 protein expression in and nearby NE cells in intrapulmonary proximal airways and trachea.**

**(A-G)** Photomicrographs of intrapulmonary proximal airways of adult *Ascl1*$^{CreERT2/+}$; *Rosa26*$^{lsl\text{-}tdTomato/+}$ lungs co-stained for Igf2 (green) and markers (red) of cell types indicated at top, with DAPI nuclear counterstain (blue). (A) NEB (NE cells, tdT$^{pos}$), (B) NE mini-cluster (NE cells, tdT$^{pos}$), (C) epithelium (E-cadherin), (D) stroma (Col1a1), (E) endothelium (Pecam1), (F) NEB-innervating pulmonary sensory neuron (PSN) (Tuj1), (G) airway smooth muscle (Acta2, also called αSMA). Note multiple Igf2$^{pos}$ NE cells in the NEB (A) and one Igf2$^{pos}$ NE cell in the mini-cluster (B). Igf2 is not detected elsewhere except apparently in airway smooth muscle (ASM, G), however ASM signal is artifactual immunolabeling or protein uptake from an external source (*139*) , as explained in fig. S6. Asterisks in (F), Igf2$^{pos}$ NEB innervated by an Igf2$^{neg}$ PSN fiber. Scale bars, 20 µm.

**(H)** Mean level and breadth of expression of Igf signaling pathway components in epithelial cell types of adult mouse trachea profiled by droplet-based scRNA-seq (*57*) (10x Genomics). Genes are rows arranged according to protein classes (receptors, transducers, binding proteins, PAPP-A proteases and inhibitors, binding protein (BP) proteases) separated by horizontal dashed lines, ordered within each class alphanumerically. Cell types are columns, with NE cells first (cyan shading) and all other epithelial cells denoted by yellow shading to indicate potential proximity to NE cells. Number of cells per type is shown below each column. Dot intensity, expression level; dot size, breadth of expression in cell type; UMI, unique molecular identifiers. Note lack of *Igf2* expression in all 96 tracheal NE cells profiled.

**(I-N)** Photomicrographs as above (A-G) in trachea. (I) NE singleton (NE cells, ZsGreen$^{pos}$, (J) epithelium (E-cadherin), (K) basal cells (Krt5), (L) stroma (Col1a1), (M) endothelium (Pecam1), (N) airway smooth muscle (Acta2). Note tracheal NE singleton (panel I) is Igf2$^{neg}$. ASM signal in (N) is again likely artifactual or due to external uptake (see fig. S6). Scale bars, 20 µm.

**(O, P)** Schematic depictions of NE and surrounding cell types in mouse intrapulmonary proximal airways (O, clustered NE cells Igf2$^{pos}$ and all surrounding cells Igf2$^{neg}$) and trachea (P, singleton NE and all surrounding cells Igf2$^{neg}$). Black dots represent Igf2 protein that is prominently but not exclusively localized at or near basal surface.

**Fig. S6**

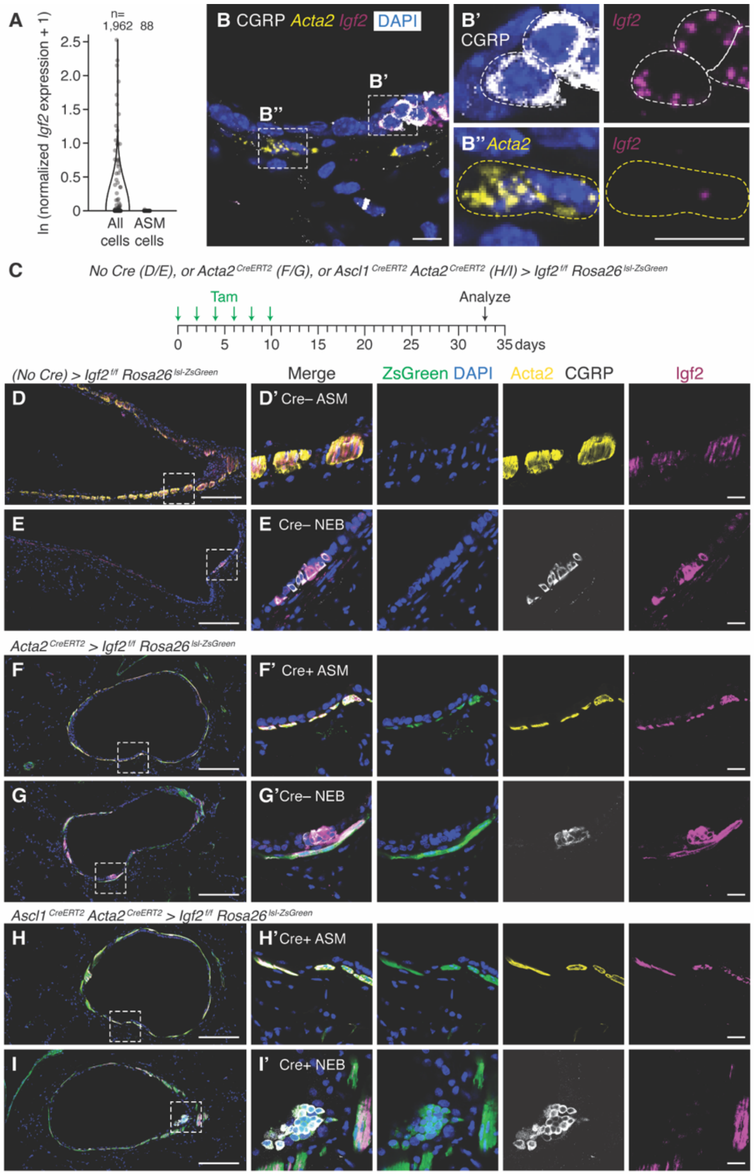


**Figure S6. Igf2 immunolabeling in airway smooth muscle cells is artifactual or represents protein uptake from an external source.**

**(A)** *Igf2* mRNA expression in mouse lung scRNA-seq profiles (M.E.K. unpublished results, see Methods). No *Igf2* is detected in airway smooth muscle (ASM) cells. fig. S3A shows similar result for additional mouse ASM cells (Ref. (*53*) ).

**(B)** Photomicrograph of NE and ASM cells from adult $Ascl1^{CreERT2/+}$; $Rosa26^{lsl\text{-}tdTomato/+}$ mice probed by smFISH (RNAscope v2) for *Igf2* mRNA (magenta) and *Acta2* mRNA (yellow, ASM), immunostained for CGRP (white, NE cells), and counterstained with DAPI (blue). (B', B") Close-ups of $CGRP^{pos}/Igf2^{pos/high}$

NE cells (B') and *Acta2*$^{pos}$/*Igf2*$^{neg/low}$ ASM (B''). Note little or no *Igf2* in ASM compared to NE cells. Scale bars, 20 µm.

**(C)** Scheme for assessing Igf2 immunostain signal in ASM. Six doses of tamoxifen (Tam) were used to induce conditional deletion of *Igf2* in adult control mice lacking a Cre driver (D, E, *Igf2*$^{flox/flox}$; *Rosa26*$^{lsl-ZsGreen}$), or with Cre drivers that delete Igf2 from ASM (*124*) (F, G, *Acta2*$^{CreERT2/+}$ ; *Igf2*$^{flox/flox}$; *Rosa26*$^{lsl-ZsGreen}$), or from ASM and NE cells (H, I, *Acta2*$^{CreERT2/+}$ ; *Ascl1*$^{CreERT2}$; *Igf2*$^{flox/flox}$; *Rosa26*$^{lsl-ZsGreen}$). Cre recombination is also marked with ZsGreen fluorophore. Analysis was done three weeks after the final induction dose to allow time for Igf2 protein turnover.

**(D-I)** Photomicrographs of airways and close-ups (boxed, insets at right) of ASM (D, F, H) and NE cells (E, G, I) of control and conditional *Igf2* deletion mutants as above co-stained for Igf2 (magenta), Acta2 (yellow, ASM), CGRP (white, NE cells), and ZsGreen (recombination marker), and counterstained with DAPI (blue). (D, E) Control mice without a Cre driver preserve *Igf2* and show no ZsGreen labeling. Note weak Igf2 signal in ASM (D, D') and strong signal in NE cells (E, E'). (F, G) ASM conditional *Igf2* deletion leaves Igf2 signal in ASM unperturbed (F, F'). (H, I) ASM and NE conditional *Igf2* deletion removes the Igf2 signal in NE cells as expected (I, I') but signal in ASM remains unperturbed (H, H'). Results demonstrate that Igf2 immunostain signal in ASM is artifactual (cross-reaction) or represents Igf2 protein uptake from an external source (*139*). Scale bars, 200 µm (D-I), 20 µm (D'-I').

**Fig. S7**

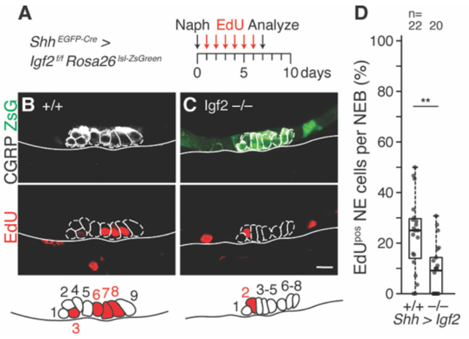

**Figure S7. Effect on NE proliferation after airway injury of conditional *Igf2* deletion from pulmonary epithelium with Shh-Cre driver.**

**(A)** Scheme for testing Igf2 requirement for NE$^{stem}$ proliferation following airway injury with naphthalene. Shh-Cre driver in *Shh$^{EGFP\text{-}Cre/+}$; Igf2$^{flox/flox}$; Rosa26$^{lsl\text{-}ZsGreen/+}$* mice is expressed early and throughout developing airway epithelium and is expected to give early and complete deletion of *Igf2* from pulmonary epithelium including NE cells. Cumulative proliferation is tracked by daily EdU injections after naphthalene airway injury.

**(B, C)** Photomicrographs and schematics of NEBs from control (B) and conditional Igf2 deletion (C) mice co-stained for CGRP (white, NE cells), ZsGreen (recombination marker), and EdU (red, proliferation marker) one week after injury. Note reduced NE proliferation after homozygous epithelial *Igf2* deletion (C). ZsGreen signal is absent in control Igf2 wild type littermate (B) due to absence of *Shh$^{EGFP\text{-}Cre}$* driver allele. Scale bars, 10 µm.

**(D)** Quantification of (A-C) showing the fraction of NE cells per NEB that proliferated by epithelial *Igf2* genotype. Note 63% proliferation reduction after homozygous deletion (Cohen's *d* -1.17, $p<10^{-2}$). n, NEBs scored.

Fig. S8

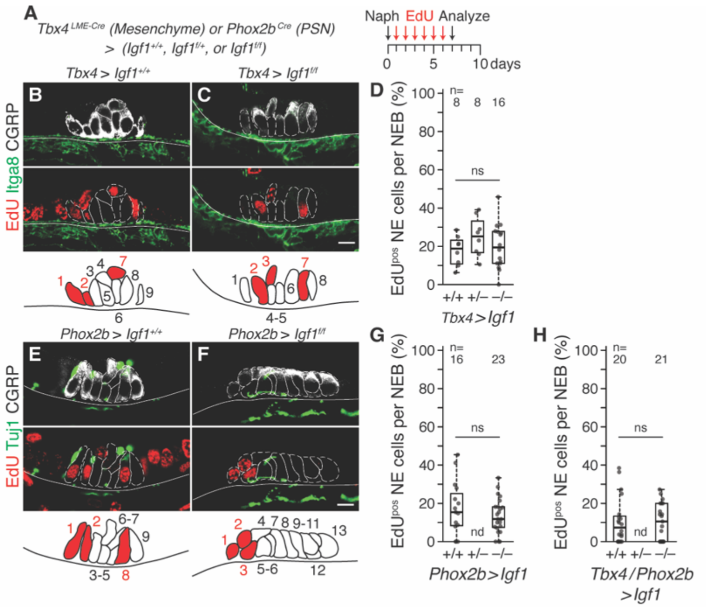

**Figure S8. Effect of stromal and neural Igf1 deletion on NE proliferation after airway injury.**

**(A)** Scheme for testing paracrine Igf1 requirement for NE$^{stem}$ proliferation following airway injury. *Tbx4$^{LME\text{-}Cre}$*; *Igf1$^{flox/flox}$* (B-D), *Phox2b$^{Cre}$*; *Igf1$^{flox/flox}$* (E-G), or *Tbx4$^{LME\text{-}Cre}$*; *Phox2b$^{Cre}$*; *Igf1$^{flox/flox}$* (H) mice were used to conditionally delete Igf1 in pulmonary mesenchyme (*Tbx4$^{LME\text{-}Cre}$* (*60*)), pulmonary sensory neurons (PSNs, *Phox2b$^{Cre}$* (*62*)), or both during embryogenesis. Daily EdU injections track cumulative proliferation for the one week interim between injury and analysis.

**(B, C)** Photomicrographs and schematics of NEBs from control wild type (B) and homozygous mesenchyme conditional *Igf1* deletion (C) adult mice co-stained for CGRP (white, NE cells), Itga8 (green, stromal cells (*140*)), and EdU (red, proliferation) one week after injury. Top row, CGRP/Itga8 merge; bottom row, Itga8/EdU merge. Note in (C) close association of at least two of the three proliferative NE cells visible (cells 2 and 7 in schematic) with Igf1 mutant stroma (Itga8$^{pos}$). Scale bars, 10 µm.

**(D)** Quantification of (A-C) showing the fraction of NE cells per NEB that proliferated by mesenchyme conditional *Igf1* deletion genotype. ns, not significant.

**(E, F)** Photomicrographs and schematics of NEBs as above but with PSN conditional *Igf1* deletion (F) and Tuj1 immunostain to detect PSN neural axons and dendrites (*141*) (green). Top row, CGRP/Tuj1 merge; bottom row, Tuj1/EdU merge. Note in (F) proliferation by at least one NE cell (cell 3 in schematic) directly contacted by an Igf1 mutant PSN dendrite (Tuj1$^{pos}$). Scale bars, 10 µm.

**(G)** Quantification of (E, F) showing the fraction of NE cells per NEB that proliferated by PSN conditional *Igf1* deletion genotype. ns, not significant.

**(H)** Quantification of NE proliferation as above following conditional *Igf1* deletion from both ASM (*Tbx4*$^{LME\text{-}Cre}$) and PSNs (*Phox2b*$^{Cre}$). ns, not significant; nd, not determined.

### Fig. S9

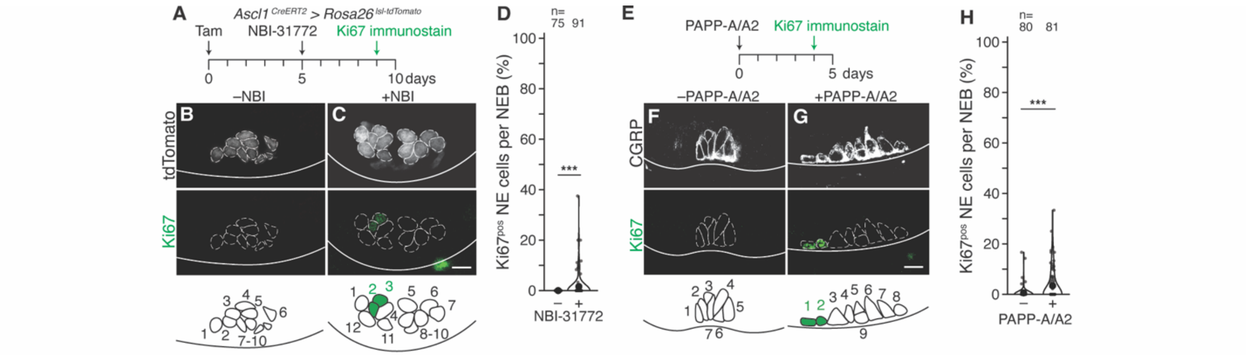


**Figure S9. Effect of pharmacologic or proteolytic disruption of Igf-Igfbp complexes on NE cell division as assessed by Ki67 immunostaining.**

**(A)** Scheme for testing the effect of acute, pharmacologic Igf-Igfbp complex dissociation on NE proliferation, as in Fig. 4F-I but with Ki67 staining instead of EdU incorporation used to assess proliferation. Tamoxifen induction of adult *Ascl1*$^{CreERT2/+}$; *Rosa26*$^{lsl\text{-}tdTomato/+}$ mice labeled NE cells (tdT$^{pos}$). Five days later, the drug NBI-31772 (240 µg) was delivered by endotracheal instillation to acutely disrupt Igf-Igfbp complexes, and then Ki67 immunostaining (*65*) was used four days later to assess active cell division. Same mice and NEBs as Fig. 4F-I.

**(B, C)** Photomicrographs and schematics of NEBs after NBI-31772 treatment for four days and co-staining for tdTomato (white, NE cells) and Ki67 (green, proliferation). Note induction of ectopic NE cell division by NBI-31772 treatment (C). Scale bars, 10 µm.

**(D)** Quantification of (A-C) showing the fraction of NE cells per NEB detected proliferating (Ki67$^{pos}$) in each condition as violin plots (mean values shown as large black dots). Note induction of NE proliferation after complex dissociation (Cohen's *d* 0.40, $p<10^{-2}$). n, number of NEB scored.

**(E)** Scheme for testing the effect of acute Igf-Igfbp complex disruption via Igfbp proteolysis to induce NE proliferation, as in Fig. 4J-M but with Ki67 staining instead of EdU incorporation used to assess proliferation. Recombinant Igfbp proteases PAPP-A (5 µg) and PAPP-A2 (5 µg) were delivered to adult C57BL/6 wild type mice by endotracheal instillation to cleave Igfbps. Ki67 immunostaining detected proliferating cells. Same mice and NEBs as Fig. 4J-M.

**(F, G)** Photomicrographs and schematics of NEBs after PAPP-A/PAPP-A2 treatment for four days and co-staining for CGRP (white, NE cells) and Ki67 (green, proliferation). Note induction of NE cell division by PAPP-A/A2 protease treatment (G). Scale bars, 10 µm.

**(M)** Quantification of (E-G) showing the fraction of NE cells per NEB detected proliferating (Ki67$^{pos}$) in each condition. Note induction of NE proliferation after PAPP-A/PAPP-A2 protease treatment (Cohen's *d* 0.57, $p<10^{-4}$). n, number of NEB scored.

**Fig. S10**

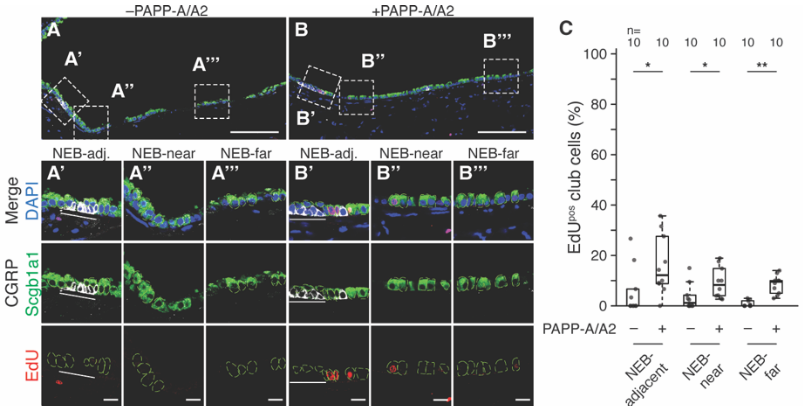


**Figure S10. Effect of proteolytic disruption of Igf-Igfbp complexes on club cell division.**

**(A, B)** Photomicrographs of airways from control and PAPP-A/A2 treated mice (same mice as Fig. 4J-M) co-stained for Scgb1a1 (club cells, green, individual cells outlined with light green), CGRP (NE cells, white), and EdU (proliferation, red), with DAPI counterstain (blue). Boxes, insets below show close-ups of club cells adjacent to NEBs ("NEB-adj.") (A', B'), near NEBs ("NEB-near") (A'', B''), and far from NEBs ("NEB-far") (A''', B'''). Note club cell proliferation adjacent to a NEB (marked by white line) after protease treatment (B'). Scale bars, 100 μm (A, B), 10 μm (insets).

**(C)** Quantification of (A, B) showing the fraction of EdU$^{pos}$ club cells scored under each condition and distance from NEBs. n, number of regions analyzed, with each data point representing the value for tens to hundreds of scored club cells. Club cell proliferation was significantly increased after drug treatment in club cells adjacent to NEBs (Cohen's *d* 1.00, $p<0.05$), and with a small effect on club cells more distant from NEBs.

**Fig. S11**

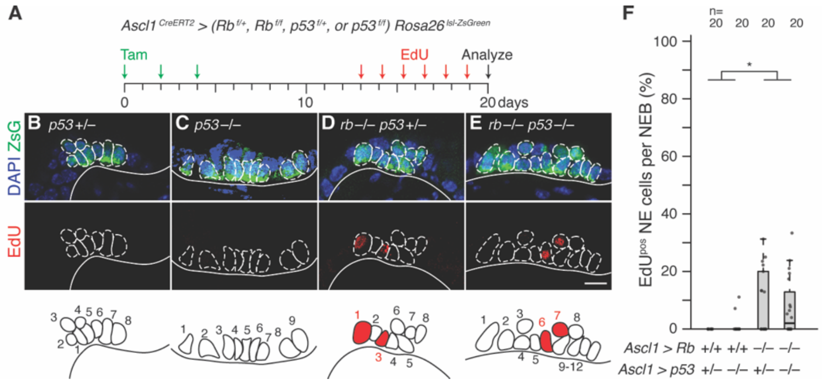


**Figure S11. Rb and p53 regulation of NE proliferation *in vivo*.**

**(A)** Scheme for testing requirements of Rb and p53 to prevent NE cell proliferation under normal (uninjured) conditions similar to (Fig. 6M) but with different genotypes. Tamoxifen induction of (B) *Ascl1*$^{CreERT2/+}$; *p53*$^{flox/+}$; *Rosa26*$^{lsl\text{-}ZsGreen/+}$, (C) *Ascl1*$^{CreERT2/+}$; *p53*$^{flox/flox}$; *Rosa26*$^{lsl\text{-}ZsGreen/+}$,(D) *Ascl1*$^{CreERT2/+}$; *Rb*$^{flox/flox}$; *p53*$^{flox/+}$; *Rosa26*$^{lsl\text{-}ZsGreen/+}$, or (E) *Ascl1*$^{CreERT2/+}$; *Rb*$^{flox/flox}$; *p53*$^{flox/flox}$; *Rosa26*$^{lsl\text{-}ZsGreen/+}$ adult mice to excise essential exons from *Rb*$^{flox}$ and *p53*$^{flox}$ and label NE cells (ZsGreen). 12 days later, daily EdU injections track proliferation over seven days.

**(B-E)** Photomicrographs and schematics showing NE cell proliferation (EdU$^{pos}$) in representative NEBs (Zsgreen$^{pos}$) of heterozygous or homozygous *p53* (B, C) or *Rb* (D, E) NE conditional deletion mutants. Note increased proliferation in homozygous *Rb* conditional deletion mutant (D, E). Scale bars, 10 µm.

**(F)** Quantification of (B-E) showing the fraction of NE cells per NEB that proliferated by NE *p53* or *Rb* NE conditional deletion genotype. n, number of NEBs scored. Note that *Rb* homozygous deletion in NE cells induced significant proliferation that matches the observed effect of *Rb* and *p53* compound deletion (Cohen's *d* -0.15, p=0.99) and increase to *p53* heterozygous (Cohen's *d* 1.01, p<0.05). By contrast, *p53* homozygous deletion alone induced little proliferation that was statistically grouped with *p53* heterozygous (Cohen's *d* 0.38, p=0.76).

**Table S1.**

**Table S1 (related to Fig. 1 and S3). Candidate mitogenic receptors queried for expression in mouse pulmonary NE cells and cognate ligands or agonists tested in P7 lung slice culture.**

| Receptor | Family | Reference | NE expression level (mean TPM) | NE expression breadth (%, n=100) | Cognate ligand or agonist tested | Working concentration (carrier solution) | NE proliferation induced (mean % EdU+ per volume) (n=volumes scored, NE cells scored) | Cohen's *d* vs. control |
|---|---|---|---|---|---|---|---|---|
| *Egfr* | RTK (Egf) | 1 | 143.2 | 49 | | | | |
| *Erbb2* | RTK (Egf) | 1 | 15.5 | 1 | Hbegf | 50 ng/ml (PBS 0.1% BSA) | 3.5 (n=8, 421) | 0.17 |
| *Erbb3* | RTK (Egf) | 1 | 21.1 | 6 | | | | |
| *Erbb4* | RTK (Egf) | 1 | 0 | 0 | | | | |
| *Fgfr1* | RTK (Fgf) | 2, 3 | 21.6 | 1 | | | | |
| *Fgfr2* | RTK (Fgf) | 2, 3 | 119.0 | 18 | Fgf1 | 100 ng/ml (PBS) | 1.4 (n=8, 532) | -0.27 |
| *Fgfr3* | RTK (Fgf) | 2, 3 | 114.7 | 1 | | | | |
| *Fgfr4* | RTK (Fgf) | 2, 3 | 0 | 0 | | | | |
| *Igf1r* | RTK (Igf) | 2, 4 | 49.0 | 50 | Igf1 | 100 ng/ml (PBS) | 21.8 (n=8, 598) | 1.75** |
| *Insr* | RTK (Igf) | 2, 4 | 62.9 | 7 | | | | |
| *Kit* | RTK | 2, 5 | 89.4 | 6 | Scf | 100 ng/ml (PBS 0.1% BSA) | 3.0 (n=7, 514) | 0.13 |
| *Ntrk1* | RTK | 6 (but see 7) | 137.4 | 1 | Ngf | 100 ng/ml (PBS) | 0.6 (n=5, 219) | -0.45 |
| *Ntrk2* | RTK | 6 | 38.7 | 25 | Bdnf | 100 ng/ml (PBS) | 0.7 (n=5, 242) | -0.42 |
| *Ntrk3* | RTK | 6 | 0 | 0 | | | | |
| *Ret* | RTK | 8 | 142.2 | 29 | Gdnf | 10 ng/ml (PBS) | 5.4 (n=8, 632) | 0.49 |
| *Ptprz1* | RTP | 9 | 195.8 | 82 | Midkine | 100 ng/ml (PBS) | 3.6 (n=2, 140) | 0.23 |
| | | | | | Pleiotrophin | 100 ng/ml (PBS) | 4.7 (n=2, 297) | 0.49 |
| *Ptch1* | Hedgehog | 10, 11 | 32.3 | 4 | | | | |
| *Ptch2* | Hedgehog | 10, 11 | 0 | 0 | SAG | 100 nM (DMSO) | 2.0 (n=8, 483) | -0.12 |
| *Smo* | Hedgehog | 10, 11 | 0 | 0 | | | | |
| *Fzd1* | Wnt | 12 | 0 | 0 | | | | |
| *Fzd2* | Wnt | 12 | 0 | 0 | | | | |
| *Fzd3* | Wnt | 12 | 81.4 | 25 | | | | |
| *Fzd4* | Wnt | 12 | 0 | 0 | | | | |
| *Fzd5* | Wnt | 12 | 72.5 | 3 | | | | |
| *Fzd6* | Wnt | 12 | 24.3 | 6 | | | | |
| *Fzd7* | Wnt | 12 | 0 | 0 | | | | |
| *Fzd8* | Wnt | 12 | 0 | 0 | | | | |
| *Fzd9* | Wnt | 12 | 0 | 0 | | | | |
| *Fzd10* | Wnt | 12 | 0 | 0 | | | | |
| *Lgr4* | Wnt | 12 | 108.8 | 12 | Wnt3a | 100 ng/ml (PBS) | 2.0 (n=4, 168) | -0.11 |
| *Lgr5* | Wnt | 12 | 64.2 | 2 | R-spondin1 (+Wnt3a) | 1 μg/ml (PBS 0.1% BSA) | 0.9 (n=4, 271) | -0.38 |
| *Lgr6* | Wnt | 12 | 0 | 0 | | | | |
| *Lrp5* | Wnt | 12 | 0 | 0 | | | | |
| *Lrp6* | Wnt | 12 | 57.6 | 26 | | | | |
| *Ptk7* | Wnt | 12 | 51.8 | 2 | | | | |
| *Rnf43* | Wnt | 12 | 151.3 | 1 | | | | |
| *Ror1* | Wnt | 12 | 0 | 0 | | | | |
| *Ror2* | Wnt | 12 | 0 | 0 | | | | |
| *Ryk* | Wnt | 12 | 92.6 | 14 | | | | |
| *Znrf3* | Wnt | 12 | 95.6 | 73 | | | | |
| *Chrna1* | nAChR | 13 | 32.9 | 49 | | | | |
| *Chrna2* | nAChR | 13 | 25.4 | 4 | | | | |
| *Chrna3* | nAChR | 13 | 0 | 0 | | | | |
| *Chrna4* | nAChR | 13 | 0 | 0 | Carbachol | 100 nM (PBS) | 4.4 (n=8, 569) | 0.46 |
| *Chrna5* | nAChR | 13 | 28.3 | 10 | | | | |
| *Chrna6* | nAChR | 13 | 0 | 0 | | | | |
| *Chrna7* | nAChR | 13 | 0 | 0 | | | | |

| | | | | | | | | |
|---|---|---|---|---|---|---|---|---|
| *Chrna9* | nAChR | 13 | 0 | 0 | | | | |
| *Chrna10* | nAChR | 13 | 11.9 | 1 | | | | |
| *Chrnb1* | nAChR | 13 | 37.8 | 10 | | | | |
| *Chrnb2* | nAChR | 13 | 55.9 | 5 | | | | |
| *Chrnb3* | nAChR | 13 | 0 | 0 | | | | |
| *Chrnb4* | nAChR | 13 | 18.0 | 3 | | | | |
| *Chrnd* | nAChR | 13 | 0 | 0 | | | | |
| *Chrne* | nAChR | 13 | 49.4 | 2 | | | | |
| *Chrng* | nAChR | 13 | 0 | 0 | | | | |
| *Ogfr* | Opioid | 13 | 47.5 | 5 | | | | |
| *Oprd1* | Opioid | 13 | 37.6 | 5 | | | | |
| *Oprk1* | Opioid | 13 | 0 | 0 | Met-enkephalin | 1 µM (PBS) | 1.3 (n=8, 540) | -0.31 |
| *Oprl1* | Opioid | 13 | 0 | 0 | | | | |
| *Oprm1* | Opioid | 13 | 0 | 0 | | | | |
| *Grpr* | Bombesin | 14 | 0 | 0 | Grp | 100 nM (water) | 2.8 (n=8, 276) | 0.07 |
| *Nmbr* | Bombesin | 14 | 0 | 0 | Nmb | 100 nM (water) | 4.9 (n=8, 420) | 0.38 |
| | | | | | Vehicle control | 0.1% BSA (PBS) | 2.5 (n=19, 1157) | |

**p=0.0095, two-sided Dunn's test with Benjamini-Hochberg multiple comparisons adjustment

| Receptor | Family | Reference | NE expression level (mean TPM) | NE expression breadth (%) | Cognate ligand or agonist tested | Concentration (vehicle) | NE proliferation induced (mean % EdU+ per volume) (n=volumes scored, NE cells scored) | Cohen's *d* vs. control |
|---|---|---|---|---|---|---|---|---|
| *Epha1* | RTK (Ephrin) | 15 | 32.7 | 3 | | | | |
| *Epha2* | RTK (Ephrin) | 15 | 177.3 | 17 | | | | |
| *Epha3* | RTK (Ephrin) | 15 | 0 | 0 | | | | |
| *Epha4* | RTK (Ephrin) | 15 | 0 | 0 | | | | |
| *Epha5* | RTK (Ephrin) | 15 | 0 | 0 | EphrinA1 (clustered) | 1 µg/ml (PBS) | 0.5 (n=4, 67) | -1.06 |
| *Epha6* | RTK (Ephrin) | 15 | 0 | 0 | | | | |
| *Epha7* | RTK (Ephrin) | 15 | 88.0 | 2 | | | | |
| *Epha8* | RTK (Ephrin) | 15 | 0 | 0 | | | | |
| *Epha10* | RTK (Ephrin) | 15 | 0 | 0 | | | | |
| *Ephb1* | RTK (Ephrin) | 15 | 0 | 0 | | | | |
| *Ephb2* | RTK (Ephrin) | 15 | 42.9 | 11 | | | | |
| *Ephb3* | RTK (Ephrin) | 15 | 83.2 | 9 | EphrinB1 (clustered) | 1 µg/ml (PBS) | 5.1 (n=4, 191) | -0.23 |
| *Ephb4* | RTK (Ephrin) | 15 | 0 | 0 | | | | |
| *Ephb6* | RTK (Ephrin) | 15 | 12.6 | 1 | | | | |
| | | | | | Ephrin control | 0.2 µg/ml (PBS) | 6.7 (n=4, 145) | |

**Table S2.**

### Table S2. Statistical Information for Figures.

| Figure | Condition | NE cells scored | NEBs scored | Mice scored | Median | Mean | Comparison | | Cohen's *d* | p-value | Test |
|---|---|---|---|---|---|---|---|---|---|---|---|
| Figure 2D | Igf1r (+/+) | 1256 | 65 | 6 | 26.5% | 28.5% | Igf1r (+/+) | Igf1r (+/–) | 0.25 | 0.26 | KWH/Dunn + BH adjustment* |
| | Igf1r (+/–) | 1798 | 55 | 5 | 23.5% | 24.8% | Igf1r (+/–) | Igf1r (–/–) | 0.65 | $1.21 \times 10^{-4}$ | |
| | Igf1r (–/–) | 1578 | 70 | 5 | 10.0% | 14.9% | Igf1r (–/–) | Igf1r (+/+) | -0.88 | $3.24 \times 10^{-7}$ | |
| Figure 2H | Igf1r (+/+) Insr (+/+) | 1148 | 52 | 3 | 20.9% | 23.4% | Igf1r (–/–) Insr (–/–) | Igf1r (+/+) Insr (+/+) | -1.62 | $5.25 \times 10^{-6}$ | KWH/Dunn + BH adjustment |
| | Igf1r (+/+) Insr (–/–) | 533 | 38 | 2 | 14.6% | 18.7% | Igf1r (+/+) Insr (+/+) | Igf1r (+/+) Insr (–/–) | -0.26 | 0.33 | |
| | Igf1r (–/–) Insr (–/–) | 907 | 69 | 4 | 0.0% | 3.0% | Igf1r (–/–) Insr (–/–) | Igf1r (+/+) Insr (–/–) | -1.14 | $7.07 \times 10^{-6}$ | |
| Figure 3A | Single (T) | 23 | 23 (singles) | 2 | 0.0% | 0.0% | Single (T) | Mini (L) | -1.38 | $3.03 \times 10^{-3}$ | KWH/Dunn + BH adjustment |
| | Mini (L) | 70 | 21 (minis) | | 50.0% | 37.4% | Mini (L) | NEB (L) | -1.19 | $2.10 \times 10^{-3}$ | |
| | NEB (L) | 266 | 17 (NEBs) | | 80.0% | 75.5% | NEB (L) | Single (T) | 6.02 | $4.31 \times 10^{-9}$ | |
| Figure 3E | WT (+/+) | 177 | 11 | 2 | 71.4% | 66.2% | Igf2 (+/+) | Igf2 (–/–) | 5.04 | $1.51 \times 10^{-5}$ | MWU |
| | KO (–/–) | 127 | 13 | 2 | 0.0% | 1.7% | | | | | |
| Figure 3F | Vehicle | 562 | 6 (volumes) | 3 | 1.4% | 2.8% | Vehicle | Igf1 | -1.27 | 0.49 | KWH/Dunn + BH adjustment |
| | Igf1 | 688 | 7 (volumes) | | 16.7% | 14.7% | | | | | |
| | Igf2 | 539 | 7 (volumes) | | 12.8% | 12.4% | Vehicle | Igf2 | -1.48 | 0.16 | |
| Figure 3J | Igf2 (+/+) | 1827 | 78 | 4 | 20.8% | 23.0% | Igf2 (+/+) | Igf2 (–/–) | 0.97 | $5.08 \times 10^{-8}$ | MWU |
| | Igf2 (–/–) | 1212 | 82 | 5 | 5.0% | 9.4% | | | | | |
| Figure 4C | Igfbp5 (+/+) | 285 | 16 | 2 | 93.1% | 89.8% | Igfbp5 (+/+) | Igfbp5 (–/–) | 4.88 | $3.20 \times 10^{-7}$ | MWU |
| | Igfbp5 (–/–) | 461 | 21 | 2 | 23.1% | 21.1% | | | | | |
| Figure 4G | Igfbp5 (+/+) | 1633 | 100 | 5 | 20.0% | 22.0% | Igfbp5 (+/+) | Igfbp5 (–/–) | -0.51 | $2.50 \times 10^{-3}$ | MWU |
| | Igfbp5 (–/–) | 1308 | 80 | 4 | 29.8% | 30.3% | | | | | |
| Figure 5D | Vehicle | 1003 | 75 | 4 | 0.0% | 0.0% | Vehicle | NBI-31772 | 0.52 | $1.45 \times 10^{-4}$ | MWU |
| | NBI-31772 | 1138 | 91 | 5 | 0.0% | 4.6% | | | | | |
| Figure 5H | Vehicle | 1269 | 80 | 4 | 0.0% | 0.0% | Vehicle | PAPP-A/A2 | -0.69 | $9.62 \times 10^{-6}$ | MWU |
| | PAPP-A/A2 | 1508 | 81 | 4 | 0.0% | 3.1% | | | | | |
| Figure 5K | Vehicle NEB adj | 109*** | 5 | 2 | 13.0% | 0.0% | Vehicle NEB adj | NBI-31772 NEB adj | -0.93 | $2.43 \times 10^{-2}$ | KWH/Dunn + BH adjustment |
| | Vehicle NEB near | 220*** | 5 | 2 | 7.0% | 0.0% | Vehicle NEB near | NBI-31772 NEB near | -0.69 | 0.14 | |
| | Vehicle NEB far | 1063*** | 5 | 2 | 6.4% | 7.0% | Vehicle NEB far | NBI-31772 NEB far | -0.95 | 0.26 | |
| | NBI-31772 NEB adj | 189*** | 5 | 2 | 33.4% | 36.9% | | | | | |
| | NBI-31772 NEB near | 456*** | 5 | 2 | 14.5% | 12.1% | | | | | |

| | | | | | | | | | | | |
|---|---|---|---|---|---|---|---|---|---|---|---|
| | NBI-31772 NEB far | 1150*** | 5 | 2 | 13.2% | 13.0% | | | | | |
| Figure 6L | Vehicle | 1391 | 15 (volumes) | 4 | 1.0% | 3.1% | Vehicle | Igf1 | -1.43 | $7.07\times 10^{-4}$ | KWH/Dunn + BH adjustment |
| | Igf1 | 1796 | 15 (volumes) | | 10.5% | 13.0% | Igf1 | Igf1 + Palbo | 1.64 | $7.19\times 10^{-4}$ | |
| | | | | | | | Igf1 | Igf1 + Nutlin-3a | 1.89 | $1.37\times 10^{-5}$ | |
| | Igf1 + Palbo | 951 | 14 (volumes) | | 1.6% | 2.2% | Igf1 + Palbo | Igf1 + Palbo + Nutlin-3 | 0.88 | 0.49 | |
| | Igf1 + Nutlin-3a | 1134 | 15 (volumes) | | 0.0% | 1.0% | Igf1 + Nutlin-3a | Igf1 + Palbo + Nutlin-3 | 0.51 | 0.17 | |
| | Igf1 + Palbo + Nutlin-3 | 820 | 13 (volumes) | | 0.0% | 0.3% | Igf1 + Palbo + Nutlin-3 | Igf1 | -1.97 | $1.21\times 10^{-6}$ | |
| | | | | | | | Vehicle | Igf2 | -1.00 | $3.53\times 10^{-2}$ | |
| | Igf2 | 1128 | 16 (volumes) | | 9.5% | 10.1% | Igf2 | Igf2 + Palbo | 1.14 | $6.67\times 10^{-4}$ | |
| | | | | | | | Igf2 | Igf2 + Nutlin-3a | 1.48 | $2.25\times 10^{-4}$ | |
| | Igf2 + Palbo | 943 | 16 (volumes) | | 0.0% | 2.0% | Igf2 + Palbo | Igf2 + Palbo + Nutlin-3 | 0.36 | 0.78 | |
| | Igf2 + Nutlin-3a | 919 | 14 (volumes) | | 0.0% | 0.5% | Igf2 + Nutlin-3a | Igf2 + Palbo + Nutlin-3 | -0.07 | 0.82 | |
| | Igf2 + Palbo + Nutlin-3 | 1195 | 12 (volumes) | | 0.0% | 0.6% | Igf2 + Palbo + Nutlin-3 | Igf2 | -1.42 | $9.85\times 10^{-4}$ | |
| Figure 6R | Rb (+/+) p53 (+/+) | 658 | 33 | 2 | 0.0% | 0.2% | Rb (+/+) p53 (+/+) | Rb (+/+) p53 (–/–) | -0.66 | 0.10 | KWH/Dunn + BH adjustment |
| | Rb (+/+) p53 (+/–) | 502 | 13 | 1 | 0.0% | 2.4% | Rb (+/+) p53 (+/–) | Rb (+/+) p53 (+/+) | 0.74 | 0.16 | |
| | | | | | | | Rb (+/+) p53 (+/–) | Rb (+/+) p53 (–/–) | 0.11 | 0.92 | |
| | Rb (+/+) p53 (–/–) | 1090 | 39 | 2 | 0.0% | 1.9% | Rb (+/+) p53 (–/–) | Rb (–/–) p53 (+/+) | -1.03 | $5.44\times 10^{-3}$ | |
| | | | | | | | Rb (+/+) p53 (–/–) | Rb (–/–) p53 (–/–) | -0.90 | $2.01\times 10^{-2}$ | |
| | Rb (+/–) p53 (+/+) | 270 | 12 | 1 | 0.0% | 1.9% | Rb (+/–) p53 (+/+) | Rb (+/+) p53 (+/+) | 0.83 | 0.28 | |
| | | | | | | | Rb (+/–) p53 (+/+) | Rb (–/–) p53 (+/+) | -0.81 | $4.47\times 10^{-2}$ | |
| | Rb (–/–) p53 (+/+) | 682 | 24 | 2 | 5.6% | 9.7% | Rb (–/–) p53 (+/+) | Rb (+/+) p53 (+/+) | 1.29 | $2.27\times 10^{-5}$ | |
| | | | | | | | Rb (–/–) p53 (+/+) | Rb (–/–) p53 (–/–) | 0.29 | 0.87 | |
| | Rb (–/–) p53 (–/–) | 490 | 23 | 2 | 5.3% | 6.9% | Rb (–/–) p53 (–/–) | Rb (+/+) p53 (+/+) | 1.35 | $1.15\times 10^{-4}$ | |
| Figure 7D | Igf1r (+/(+/–)) Insr (+/(+/–)) | NA | 538 | 5 | 27.1% | 22.8% | Igf1r (+/(+/–)) Insr (+/(+/–)) | Igf1r (–/–) Insr (–/–) | 2.01 | $1.60\times 10^{-2}$ | MWU |
| | Igf1r (–/–) Insr (–/–) | NA | 333 | 5 | 8.3% | 8.5% | | | | | |
| Figure 7E | Igf1r (+/(+/–)) Insr (+/(+/–)) | NA | 127 | 5 | 10.7% | 19.6% | Igf1r (+/(+/–)) Insr (+/(+/–)) | Igf1r (–/–) Insr (–/–) | -0.28 | 0.45 | MWU |
| | Igf1r (–/–) Insr (–/–) | NA | 29 | 5 | 15.3% | 25.6% | | | | | |
| Figure S1E | Egfr (+/+) | 600 | 22 | 1 | 27.8% | 29.1% | Egfr (+/+) | Egfr (+/–) | 0.55 | 0.48 | KWH/Dunn + BH adjustment |
| | Egfr (+/–) | 304 | 14 | 2 | 22.2% | 21.5% | Egfr (+/–) | Egfr (–/–) | -0.44 | 0.50 | |
| | Egfr (–/–) | 458 | 15 | 4 | 31.3% | 27.9% | Egfr (–/–) | Egfr (+/+) | -0.09 | 0.72 | |
| Figure S4D | Vehicle *Igf1r*$^{neg}$ | 203 | 20 | 2 | N.A. | 0.0% | Naph EdU$^{pos}$ *Igf1r*$^{neg}$ | Naph EdU$^{pos}$ *Igf1r*$^{pos}$ | N.A. | $4.72\times 10^{-2}$ | Fisher's exact test |
| | Vehicle *Igf1r*$^{pos}$ | 130 | 20 | 2 | N.A. | 0.0% | | | | | |
| | Naph *Igf1r*$^{neg}$ | 82 | 10 | 2 | N.A. | 25.6% | | | | | |
| | Naph *Igf1r*$^{pos}$ | 61 | 10 | 2 | N.A. | 31.7% | | | | | |

| | | | | | | | | | | | |
|---|---|---|---|---|---|---|---|---|---|---|---|
| Figure S7D | Igf2 (+/+) | 341 | 20 | 1 | 25.0% | 23.3% | Igf2 (+/+) | Igf2 (–/–) | 1.17 | $1.21 \times 10^{-3}$ | MWU |
| | Igf2 (–/–) | 288 | 22 | 1 | 9.2% | 8.9% | | | | | |
| Figure S8D | Igf1 (+/+) | 215 | 8 | 1 | 18.9% | 17.6% | Igf1 (+/+) | Igf1 (+/–) | -0.82 | 0.50 | KWH/Dunn + BH adjustment |
| | Igf1 (+/–) | 263 | 8 | 1 | 25.2% | 25.0% | Igf1 (+/–) | Igf1 (–/–) | 0.44 | 0.44 | |
| | Igf1 (–/–) | 454 | 16 | 3 | 19.4% | 20.0% | Igf1 (–/–) | Igf1 (+/+) | 0.23 | 0.58 | |
| Figure S8G | Igf1 (+/+) | 435 | 16 | 2 | 15.1% | 16.3% | Igf1 (+/+) | Igf1 (–/–) | 0.25 | 0.71 | MWU |
| | Igf1 (–/–) | 740 | 23 | 3 | 11.8% | 13.5% | | | | | |
| Figure S8H | Igf1 (+/+) | 406 | 20 | 1 | 7.3% | 10.6% | Igf1 (+/+) | Igf1 (–/–) | -0.03 | 0.89 | MWU |
| | Igf1 (–/–) | 449 | 21 | 1 | 10.5% | 10.3% | | | | | |
| Figure S10D | Vehicle | 1003 | 75 | 4 | 0.0% | 0.0% | Vehicle | NBI-31772 | -0.40 | $1.93 \times 10^{-3}$ | MWU |
| | NBI-31772 | 1138 | 91 | 5 | 0.0% | 1.6% | | | | | |
| Figure S10H | Vehicle | 1269 | 80 | 4 | 0.0% | 0.6% | Vehicle | PAPP-A/A2 | -0.57 | $5.95 \times 10^{-5}$ | MWU |
| | PAPP-A/A2 | 1508 | 81 | 4 | 0.0% | 3.4% | | | | | |
| Figure S11C | Vehicle NEB adj | 247*** | 5 | 2 | 5.5% | 0.0% | Vehicle NEB adj | PAPP-A/A2 NEB adj | -1.00 | $1.57 \times 10^{-2}$ | KWH/Dunn + BH adjustment |
| | Vehicle NEB near | 457*** | 5 | 2 | 3.4% | 1.2% | Vehicle NEB near | PAPP-A/A2 NEB near | -1.02 | 0.05 | |
| | Vehicle NEB far | 1400*** | 5 | 2 | 0.8% | 0.0% | Vehicle NEB far | PAPP-A/A2 NEB far | -2.81 | $8.36 \times 10^{-3}$ | |
| | PAPP-A/A2 NEB adj | 201*** | 5 | 2 | 16.1% | 12.1% | | | | | |
| | PAPP-A/A2 NEB near | 343*** | 5 | 2 | 9.2% | 8.3% | | | | | |
| | PAPP-A/A2 NEB far | 1200*** | 5 | 2 | 8.5% | 9.5% | | | | | |
| Figure 12F | Rb (+/+) p53 (+/–) | 133 | 11 | 1 | 0.0% | 0.0% | Rb (+/+) p53 (+/–) | Rb (–/–) p53 (+/–) | -1.01 | $1.31 \times 10^{-2}$ | KWH/Dunn + BH adjustment |
| | Rb (+/+) p53 (–/–) | 316 | 21 | 1 | 0.0% | 0.9% | Rb (+/+) p53 (+/–) | Rb (+/+) p53 (–/–) | -0.38 | 0.76 | |
| | | | | | | | Rb (–/–) p53 (+/–) | Rb (+/+) p53 (–/–) | 1.03 | $1.06 \times 10^{-2}$ | |
| | Rb (–/–) p53 (+/–) | 219 | 18 | 1 | 0.0% | 9.0% | Rb (–/–) p53 (+/–) | Rb (–/–) p53 (–/–) | 0.15 | 0.99 | |
| | | | | | | | Rb (+/+) p53 (+/–) | Rb (–/–) p53 (–/–) | -0.90 | $3.49 \times 10^{-2}$ | |
| | Rb (–/–) p53 (–/–) | 410 | 20 | 1 | 0.2% | 7.3% | Rb (+/+) p53 (–/–) | Rb (–/–) p53 (–/–) | -0.88 | $1.79 \times 10^{-2}$ | |

* KWH/Dunn + BH adjustment: nonparametric Kruskal-Wallis H tests, using the kruskal.test function (stats), followed by Dunn's test to explicitly compare distributions pairwise, using the function dunnTest (FSA)
** MWU: nonparametric Mann-Whitney U test (also known as the Wilcoxon rank-sum test), implemented using the wilcox.test function from the base stats package
*** NE cells scores refers to club cells scores
